# HDAC inhibitors decrease TLR7/9-mediated human plasmacytoid dendritic cell activation by interfering with IRF-7 and NF-κB signaling

**DOI:** 10.1101/2020.05.09.085456

**Authors:** Dante B. Descalzi-Montoya, Jihong Dai, Sukhwinder Singh, Patricia Fitzgerald-Bocarsly

## Abstract

Histone deacetylase inhibitors (HDACi) are epigenome modulating molecules that target histone and non-histone proteins and have been successfully used to target many types of cancer and immunological disorders. While HDACi’s effects on nuclear histone deacetylases are well characterized, their effect on non-nuclear, cytoplasmic molecules requires further investigation. In the current study we characterized the effects of class I/II HDACi, specifically, TSA, MS-275, and SAHA, on plasmacytoid dendritic cell (pDC) biology upon viral activation via the TLR7/9 pathway. TSA, MS-275, and SAHA, down-modulated the induction of IFN-α and TNF-α upon Influenza A virus (IAV; TLR7 signaling) and Herpes Simplex 1 (HSV-1; TLR9 signaling) stimulation in primary pDC. The HDACi inhibitory effect was more prominent for IAV-mediated responses than for HSV-1. While IFN-α induction was not associated with inhibition of IRF-7 upregulation in the presence of TSA or MS-275, IRF-7 upregulation was affected by SAHA, but only for IAV. Furthermore, TSA, but not MS-275, inhibited TLR7/9-induced expression of maturation markers, CD40, and CD86, but not CD40. In addition, HDACi treatment increased virally-induced shedding of CD62L. Mechanistically, TSA, MS-275, SAHA significantly decreased early IRF-7 and NF-κB nuclear translocation, which was preceded by a decline in phosphorylation of IRF-7 at Ser477/479 and NF-κB p-p65, except for MS-275. In summary, we propose that broad HDACi, but not class I HDACi, treatment can negatively impact early TLR7/9-mediated signaling, namely, the disruption of IRF-7 and NF-κB activation and translocation that lead to deleterious effects on pDC function.

## Introduction

Plasmacytoid dendritic cells (pDC) are a potent sub-population of dendritic cells that importantly bridge the innate and adaptive immune systems. They have the ability to rapidly produce high amounts of Type I and Type III interferon (IFN) upon challenge with synthetic TLR7 and TLR9 agonists or with DNA or RNA viruses, even in the absence of viral gene expression (reviewed in [1, 2]). TLR activation induces MyD88-dependent downstream signaling that subsequently leads to the phosphorylation and nuclear translocation of Interferon Regulatory Factor-7 (IRF-7), which pDC also express constitutively at a basal level, resulting in the expression of type I and III interferons [2–5].In addition, pDC TLR activation can also induce production of pro-inflammatory cytokines such as TNF-α and IL-6, and chemokines CXCL10 (IP-10) and CCL5 (RANTES) [6]. In addition, pDC have the ability to migrate from peripheral blood to peripheral lymphoid tissue and sites of inflammation via their constitutive expression of CD62L [7]. IFN-α production in pDC can be triggered by viral or synthetic stimuli, including viral DNA or non-methylated CpG oligonucleotides, respectively, which activate endosomal Toll-like receptor (TLR)-9, or viral single-stranded RNA, or small molecules of the imidazoquinoline family, both of which trigger endosomal TLR-7.

The gene expression of Interferons is partially regulated by post-translational modifications of the *ifn* promoter and specific enzyme complexes known as histone deacetylases (HDAC) [8]. HDAC and histone acetyl transferases are enzymes that work together to regulate the acetylation state of histone and non-histone intracellular substrates [9]. They exert their effects by deacetylating ε-acetyl-lysine residues on the amino-terminal end of histone tails and non-histone targets residues. There are four HDAC classes currently characterized based on their structural similarity to homologous proteins in *Saccharomyces cerevisiae*, two of which are relevant to this manuscript: Class I and II HDACs depend on Zn^++^ for their activity. Class I HDACs include 1, 2, 3 and 8 and class II HDACs include class IIa (4, 5, 7, and 9), and class IIb (6 and 10).

Most of the intricate workings of HDAC biology have been elucidated with the utilization of HDAC inhibitors (HDACI), which are small naturally or artificially made molecules that have demonstrated great potential for the treatment of tumors *in vitro* and *in vivo*, as cancer cells are more sensitive to HDACI treatment than healthy cells. HDACI induce apoptosis in cancer cells by releasing the cell cycle block that truncates healthy terminal cell differentiation.

The Class I HDAC complexes that have been extensively studied are nuclear SIN3, NURD, N-CoR, and SMRT, which have important roles in gene regulation, including deacetylation, binding of histone proteins, ATP-dependent nucleosome remodeling, and chromosome scaffolding. cDNA array studies have shown that only up to 10% of genes are up- or down-regulated in myeloma, colon carcinoma, and leukemia cell lines when treated with several HDACi including Tricostatin A (TSA), MS-275, and Vorinostat (Suberoyanilide Hydroxamic Acid, SAHA) [10, 11]. Because of this very low effect on gene expression via modification of histone complexes, it had been suggested that the cellular effects exerted by HDACI must also occur as a result of their effects on non-histone targets. Indeed, the acetylation-dependent signal transduction observed for the type I IFN receptor (IFNAR1/2) is a very good example. Tang *et al*. showed that CREB-binding protein acetylates the SH2 cytoplasmic domain of IFNAR2 causing recruitment of IRF-9 and subsequent formation of the ISGF3 complex, which are necessary for signal transduction [12]. Another example is the observed effect that inhibition of HDAC6 by HDACi or siRNA methods dramatically increases the acetylation of α-tubulin and its association with cortactin, which results in dramatic cytoskeletal changes [13, 14].

TSA and SAHA have been shown to be quite effective in decreasing immunological disorders in animal models. TSA reduced leukocyte infiltration and cytokine production in murine models of airway inflammation, colitis, arthritis, and transplant rejection [15, 16]. TSA also has been shown to down modulate the expression of inflammatory genes upon TLR engagement in murine macrophages and dendritic cells, as well as inhibit T cell activation [17–19]. Particularly, in pDC TSA has been shown to inhibit the production and release of IFN-α, TNF-α, and IL-6 upon CpG stimulation by affecting IFN mRNA steady state levels [20, 21]. SAHA also has been implicated in the downregulation of immune responses. For example, SAHA reduced LPS- or IFN-γ-induced IL-12 and IL-18 production from human PBMC and LPS-induced IL-12 and Edn-1 mRNA levels from murine bone-marrow derived monocytes [18, 22].

In this study, we report the potential of TSA, MS-275and SAHA for partially interfering with virus-induced human pDC IFN-α and TNF-α production, maturation and activation by disturbing TLR7/9-mediated signal transduction events *in vitro*. We propose that the early regulation of phosphorylation of cytoplasmic transcription factors can partially explain the inhibition of pDC function in the context of viral activation.

## Materials & Methods

### Cell preparation

Heparinized peripheral blood was obtained from consenting healthy adult donors under a protocol approved by the Rutgers NJMS Institutional Review Board. PBMC were isolated using Ficoll-hystopaque (Sigma-Aldrich) gradient centrifugation and resuspended at 1-2 x 10^6^ cells/ml in RPMI-10% FCS (RPMI 1640 w/L-glutamine (Cellgro Gibco) supplemented with 10% FCS (Cellgro Gibco), 1% PEN/STREP (Cellgro Gibco), 25 mM HEPES buffer (Sigma-Aldrich). For some experiments PBMC were further enriched for pDC using negative isolation kit (Pan-DC Enrichment kit, human). H9 T cells were obtained through the NIH AIDS Reagent Program, Division of AIDS, NIAID, and NIH maintained in RPMI-10% FCS.

### TLR agonists

Herpes simplex virus type 1 (HSV-1) strain 2931 (originally obtained from Dr. Carlos Lopez, then at the Memorial Sloan-Kettering Cancer Center, New York, USA) was grown and tittered in VERO cells. HSV-1 was used to stimulate PBMC or pDC at an MOI of 1 and Influenza A Virus (IAV) strain PR8 (Charles River Laboratories, Wilmington Massachusetts, USA) was used at an MOI of 2. For intracellular cytokine detection, 5 μg/ml of Brefeldin A (Sigma-Aldrich) was added 2 hours prior to the end of the incubation period. CpG-B ODN #1826 (Invivogen) was used at 5 μg/ml for stimulation in all experiments.

### HDAC inhibitors (HDACi)

Trichostatin A (TSA) was purchased from Sigma-Aldrich (St. Louis, MO) and used in a dose response or at 330 nM (100 ng/ml) fixed concentration; MS-275 was used at 5 μM (Selleck Chemicals, Houston, TX). Suberoylanilide hydroxamic acid (SAHA, Vorinostat) used in a dose response or at a 500 nM fixed concentration (Sigma-Aldrich, St. Louis, MO). The vehicle control

DMSO(Sigma-Aldrich, St. Louis, MO) and was used at 0.01% to match TSA and SAHA concentrations, 1% to match MS-275, and 5% to match MC1568.

### pDC maturation assay

PBMC were stimulated with HSV-1, IAV, or CpG-B ODN (5 μg/ml) for 8 hours and then surface staining was performed using the following abs anti-human BDCA-2-PE (clone AC144; Biolegend), CD123-PeCy7 (clone 6H6; Biolegend), CD80 PerCPCy5.5 (clone 2D10; Biolegend), -CD40 APCCy7 (clone SC3; Biolegend), and -CD62L AF700 (DREG-56; Biolegend), CD86 FITC (clone 2331 (FUN-1); BD Biosciences). Isotype controls were used at the same concentrations as the corresponding antibodies and were the following: PerCPCy5.5 IgG1 (MOPC-21; Biolegend), APCCy7 IgG1 (MOPC-21; Biolegend), AF700, IgG1 (MOPC-21; Biolegend), FITC IgG1 (MOPC-21; BD Biosciences), κ isotype controls.

### IFN-α ELISA

PBMC were activated with TLR7/9 stimuli for 18 hr. in the presence or absence of HDACI and supernatants were collected and stored at −80° C. Supernatants were then thawed and assayed using the human IFN-α module set for ELISA (Bender MedSystems). Absorbance was measured at 450 nm using a GENios TECAN ELISA plate reader.

### Surface and Intracellular Staining

For surface staining, cells were washed with cold 0.1% BSA in PBS and blocked with 5 μl of heat-inactivated, pooled human serum for 5 minutes. Surface antibody combinations were used to specifically identify pDC, anti-human BDCA-2-PE (clone AC144; Miltenyi Biotech) and anti-human CD123-APC (clone 6H6; Biolegend). Cells were incubated with surface antibodies at 4°C for 30 minutes, then washed once with cold wash buffer, and subsequently fixed in 1% PBS overnight; if intracellular staining was to follow the next day. For intracellular staining overnight-fixed samples were washed with 2% FCS in PBS and then permeabilized with either 0.1% Triton-X (for intracellular cytokine and transcription factor staining) in 2% FCS in PBS for 5 minutes or 0.5% Saponin (for intracellular cytokine staining) in PBS for 15 minutes at room temperature (RT). Cells were washed and incubated with antibodies to human IFN-α biotin (clone MMHA2; PBL), TNF-α-Pacific Blue (clone MAb11; Biolegend) or TNF-α FITC (clone cA2; Miltenyi Biotech), and Zenon-labeled (Invitrogen)-anti-IRF-7-AF488 (clone, G-8, Santa Cruz) or AF488 Zenon-labeled (Invitrogen) mouse IgG2a isotype control (clone UPC-10; Sigma-Aldrich,), for 30 minutes at RT. Then, samples were washed with PBS once and then stained with Streptavidin-PECY7 (Biolegend) for 30 minutes at RT. Samples were then washed twice, once with wash buffer and then with PBS, and then samples were fixed in 1% PFA in PBS. Samples were processed in a BD LSR II flow cytometer by acquiring 300,000 events, then data were analyzed with FlowJo software 6.0v.

### Nuclear translocation assay

Enriched pDC were stimulated for 4 hr with HSV-1 (MOI of 1) or IAV (MOI of 2) in the presence or absence of TSA, SAHA, or DMSO vehicle. Samples were then surface-stained using anti-human BDCA-2-PE (clone AC144; Miltenyi Biotech). After the cells were washed and fixed overnight, they were washed with 2% FCS in PBS and permeabilized with 0.1% Triton-X in 2% FCS in PBS for detection of human IRF-7 or with 0.5% Triton-X 2% FCS in PBS for detection of human NF-κB, for 5 min. at RT. In separate experiments, cells were washed and directly stained with Zenon-labeled anti-human IRF-7-AF488 (clone G-8; Santa Cruz) or indirectly stained with purified anti-NF-kB p65 antibody (clone poly6226; Biolegend) for 30 minutes. Samples were then washed with 2% and stained with 2% FCS in PBS and subsequently washed stained with goat anti-rabbit IgG –FITC secondary (BD) for 30 minutes at RT. Samples were then washed with 2% FCS in PBS and transferred to 0.5 ml low adhesion Eppendorf tubes for acquisition by ImageStream IS100. DRAQ5 (Invitrogen) was used at 1:500 to stain the nuclei prior to acquisition and 10,000 events were acquired from each sample. Data were analyzed using IDEAS software 5.0v. First, single events were identified by plotting the Bright Field (BF) aspect ratio vs. area, then events in focus were gated in a gradient RMS histogram of the DRAQ5 nuclear stain, and IRF-7^+^/BDCA-2^+^ pDC were gated on a dot plot of the BDAC-2 vs. IRF-7 intensities, finally the percent of IRF-7 translocation was gated in a IRF-7/DRAQ5 similarity score histogram and reported as % translocated.

### BD Phosflow^™^ Assay

Phosphorylation of IRF-7 and NF-κB p65 was measured at 3 hr after HSV-1 or IAV stimulation by adding 1 ml of BD Cytofix to 1 ml of cell culture for 10 minutes at 37°C, then cells were washed with 2% FCS-PBS and surface-stained with anti-human HLA-DR-Pacific Blue or -APC (clone L243; Biolegend) and anti-human CD123-FITC or -PeCy5 (clone 6H6; Biolegend) antibodies for 30 minutes at 4°C. Cells were washed and fixed with 300 μl of 2% PFA - PBS on ice for 10 minutes. Then samples were permeabilized on ice by adding, drop by drop, cold BD Perm Buffer III for detection of phosphorylated IRF-7 (pIRF-7); or cold 70% ethanol, for detection of phosphorylated NF-κB p65 (pNF-κB). Samples were incubated on ice for 1 hr., then washed with 2% FCS - PBS and stained with Mouse anti-IRF-7-PE (pS477/pS479, clone K47-671; BD Biosciences) or Mouse anti-NF-κB p65-Alexa Fluor 647 (pS529, clone k10-895.12.50; BD Biosciences) or for 30 min. at RT. Cells were subsequently washed and fixed with 1% PFA-PBS for flow cytometric analysis. PBMC were gated based on SSC-A vs. FSC-A (Area) and pDC were identified by gating on double positive HLA-DR^high^/CD123^high^ events on a HLA-DR-APC vs. CD123-PE intensity dot plot, then the percentage of phosphorylated IRF-7^+^ or NF-κB p65^+^ was identified based on the unstimulated control.

### Statistics

Statistical analyses were performed using GraphPad Prism software 5.0. Data are expressed as mean ± SEM; Data were analyzed with one-way ANOVA with Bonferroni’s post hoc test; * = p<0.05; ** = p<0.01; *** = p<0.001).

## Results

### HDACi inhibitHSV-1 andIAV-inducedpDCproduction of?FN-a and TNF-α and upregulation of maturation markers

Several studies have shown the effects of HDACi on the function of dendritic cells, macrophages, and T lymphocytes [17, 18, 20, 23–25], mostly in the context of mitogen treatment, such as phytohemagglutinin or concanavalin A for T cells or synthetic TLR stimulants, such as, lipopolysaccharide or CpGs. This study focuses on the effects caused by HDACI on human pDC biology while using natural TLR7/9 viral stimulators that mimic natural viral activation. We pre-treated human PBMC with broad class I/II HDACi TSA for 1 hr., and included MS-275 in order to compare the inhibition profile when only a class I HDACi is used, then stimulated with either TLR9-specific viral stimulant HSV-1 or TLR7-specific viral stimulant IAV for 6 hrs. After incubation, PBMC were processed for surface staining for specific pDC markers and intracellular staining for IFN-α and TNF-α production. A representative analysis from dosing experiments, demonstrating the effects of TSA and MS-275 on HSV-1 and IAV-induced cytokine production of IFN-α and TNF-α at 6 hours, is shown in **Fig. 1A and 1C**, along the gating strategy utilized to derive the data **(Suppl. Fig. 1)**. In these experiments a dose-dependent inhibition of intracellular IFN-α and TNF-α that reached statistical significance at 100 ng/ml (330 nM) for TSA **(Fig. 1B)** and 5 μM for MS-275 **(Fig. 1D)** was observed, except for TNF-α, for which only a negative trend occurred with MS-275. Our findings were consistent with observations by Salvi *et al*. that used synthetic TLR agonists, but not natural ligands [20]. Since, the concentrations were found to be non-toxic for pDC by Annexin-V/7-AAD staining and did not affect pDC numbers at 6 hours **(Suppl. Fig. 2),** they were used in subsequent experiments. Furthermore, the inhibitory effect by TSA **(Fig. 2A)** and MS-275 **(Fig. 2B)** persisted at 18 hours in PBMC supernatants by ELISA, for which a more prominent inhibition was observed when using IAV, a TLR7 stimulant, than with HSV-1, which signals for IFN production through TLR9.

**FIGURE 1.**
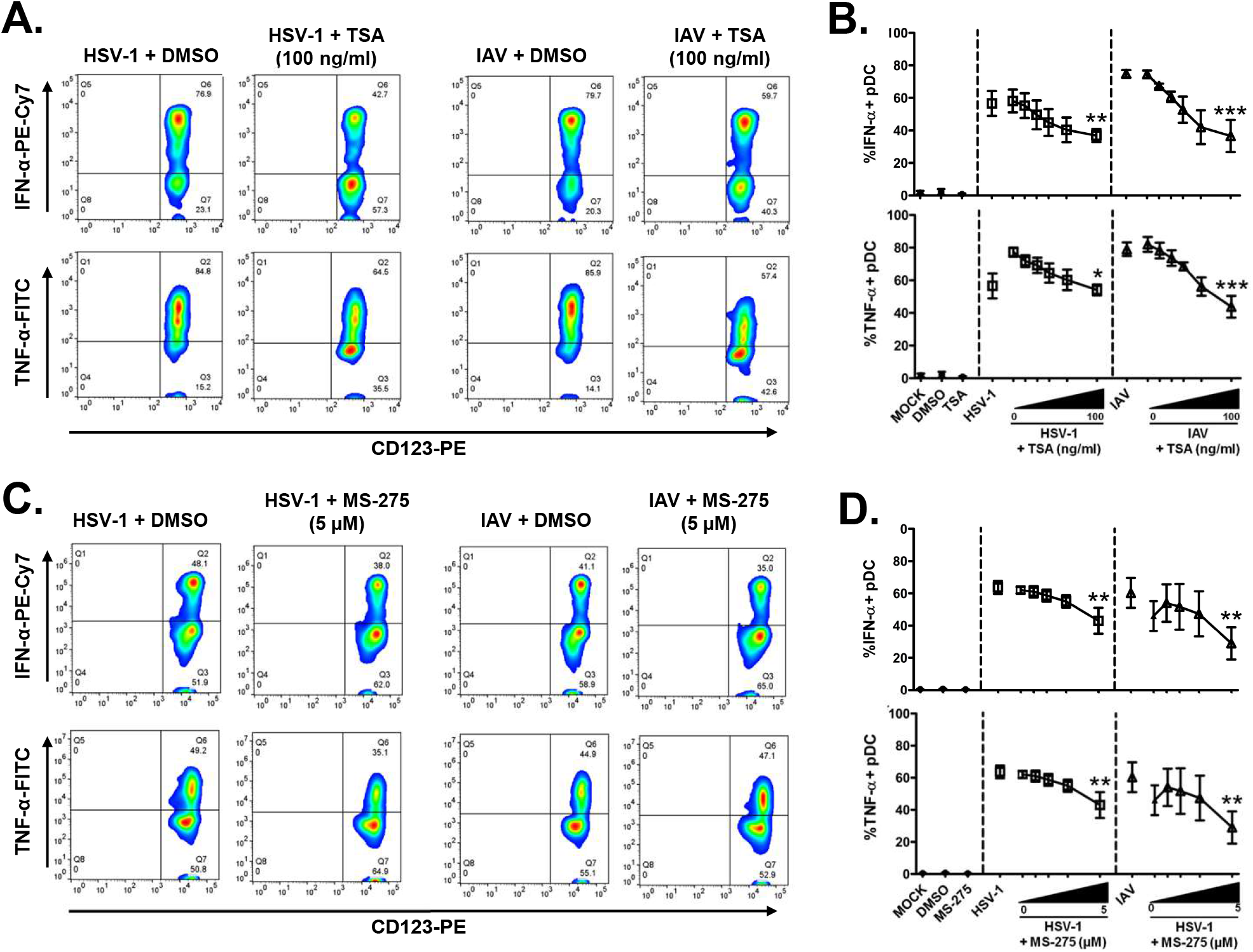
TSA and MS-275 inhibit HSV-1 and IAV-induced IFN-α and TNF-α production in human pDC in a dose-dependent manner. PBMC were incubated with TSA and MS-275 at decreasing concentrations for 1 hour and then stimulated with HSV-1 or IAV for 6 hours. Cells were then processed for intracellular flow cytometry to detect IFN-α and TNF-α production in pDC. Representative histograms showing the differential inhibition of intracellular HSV-1 and IAV-induced IFN-α and TNF-α production in pDC by TSA (100 ng/ml = 330 nM) or DMSO vehicle control (1:1000) **(A),** and MS-275 (5 μM) or DMSO vehicle control (1:530) **(C).** Pooled data of dose curve experiments showing a dose-dependent inhibitory effect on HSV-1- (MOI of 1; squares) or IAV (MOI of 2; triangles) -induced IFN-α and TNF-α by TSA (0-100 ng/ml) **(C)** and MS-275 (0-5 μM) **(D)** when compared to their respective vehicle controls. (TSA, N=3; MS-275, N=4; 3-4 independent experiments from different donors; Data are expressed as mean ± SEM; Data were analyzed with 1-way ANOVA with Bonferroni’s post test; *p<0.05; **p<0.01; ***p<0.001).

**FIGURE 2.**
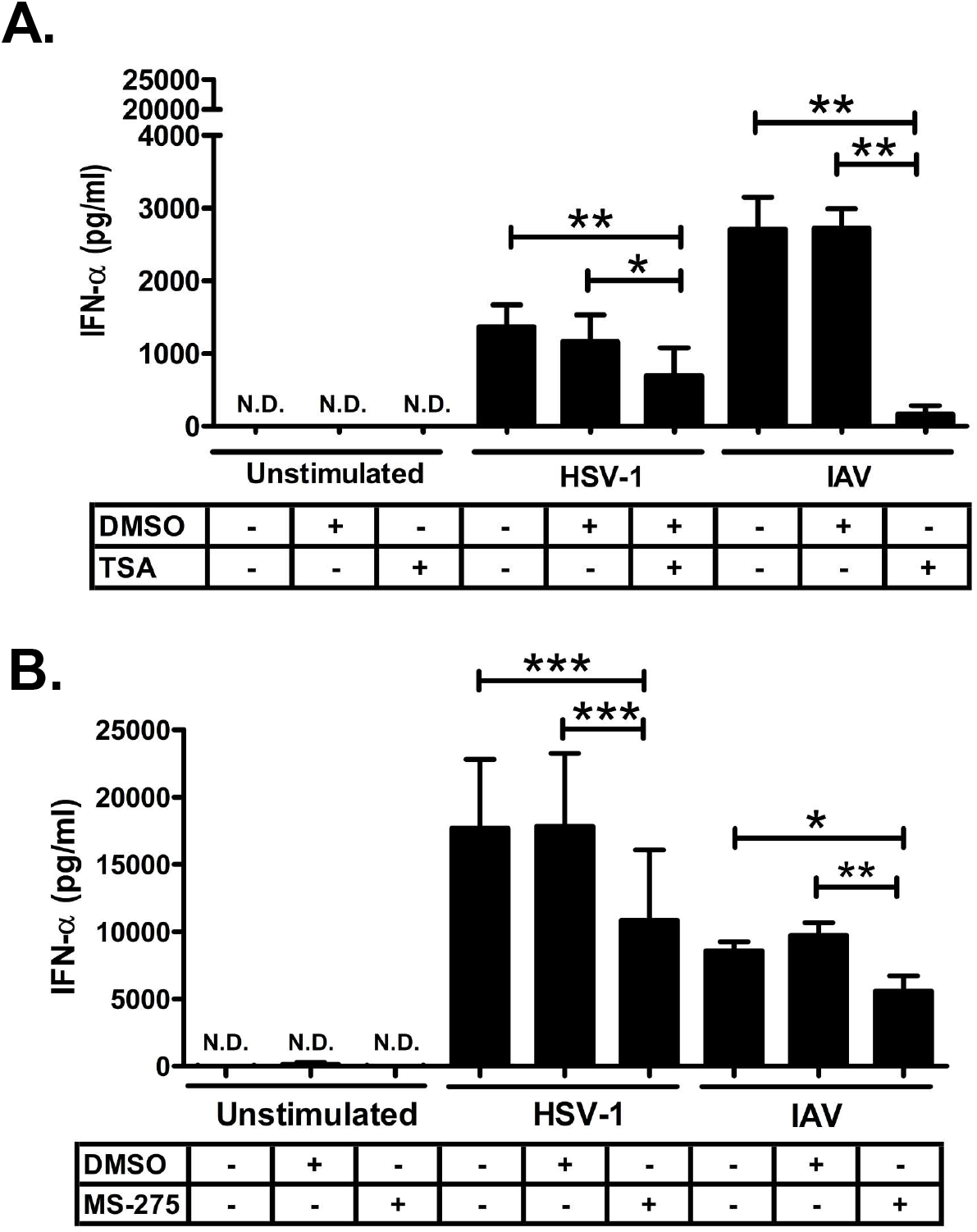
TSA and MS-275 inhibit HSV-1 and IAV-induced total IFN-α production in human pDC. PBMC were isolated and pre-treated with TSA (100 ng/ml) or DMSO vehicle control (1:1000) **(A)** and MS-275 (5 μM) or DMSO vehicle control (1:530) **(B)** for 1 hour and then stimulated with HSV-1 (MOI of 1) and IAV (MOI of 2) for 18 hours. Supernatants were collected and tested for IFN-α production by an ELISA assay. (N=4, 4 independent experiments from different donors; Data are expressed as mean ± SEM; Data were analyzed with 1-way ANOVA with Bonferroni’s post test; *p<0.05; **p<0.01; ***p<0.001; N.D.= not detected)

The effects by HDACi on TLR-induced pDC maturation were differential. TSA, but not MS-275, significantly and consistently inhibited CD86 and CD40 upregulation in pDC **(Fig. 3A-C)**. Interestingly, the upregulation of CD83 expression was inhibited by TSA only when IAV was the stimulus, but not for HSV-1 or CpG-B, which are TLR9 agonists **(Fig. 3C)**. Additionally, there was no effect on CD40 upregulation by MS-275 for both TLR7 and 9stimulation **(Fig. 2C).** Moreover, the presence of the homing molecule CD62L (L-selectin), which is shed by pDC upon activation by viruses, was decreased by TSA only with HSV-1 or CpG-B TLR9 engagement, but not with IAV, whereas MS-275 had no effect **(Fig. 2E).** These effects on maturation markers were associated with inhibition of IFN-α at 6 hours, which verified the previously demonstrated inhibitory effect by TSA and MS-275 **(data not shown).**

**FIGURE 3.**
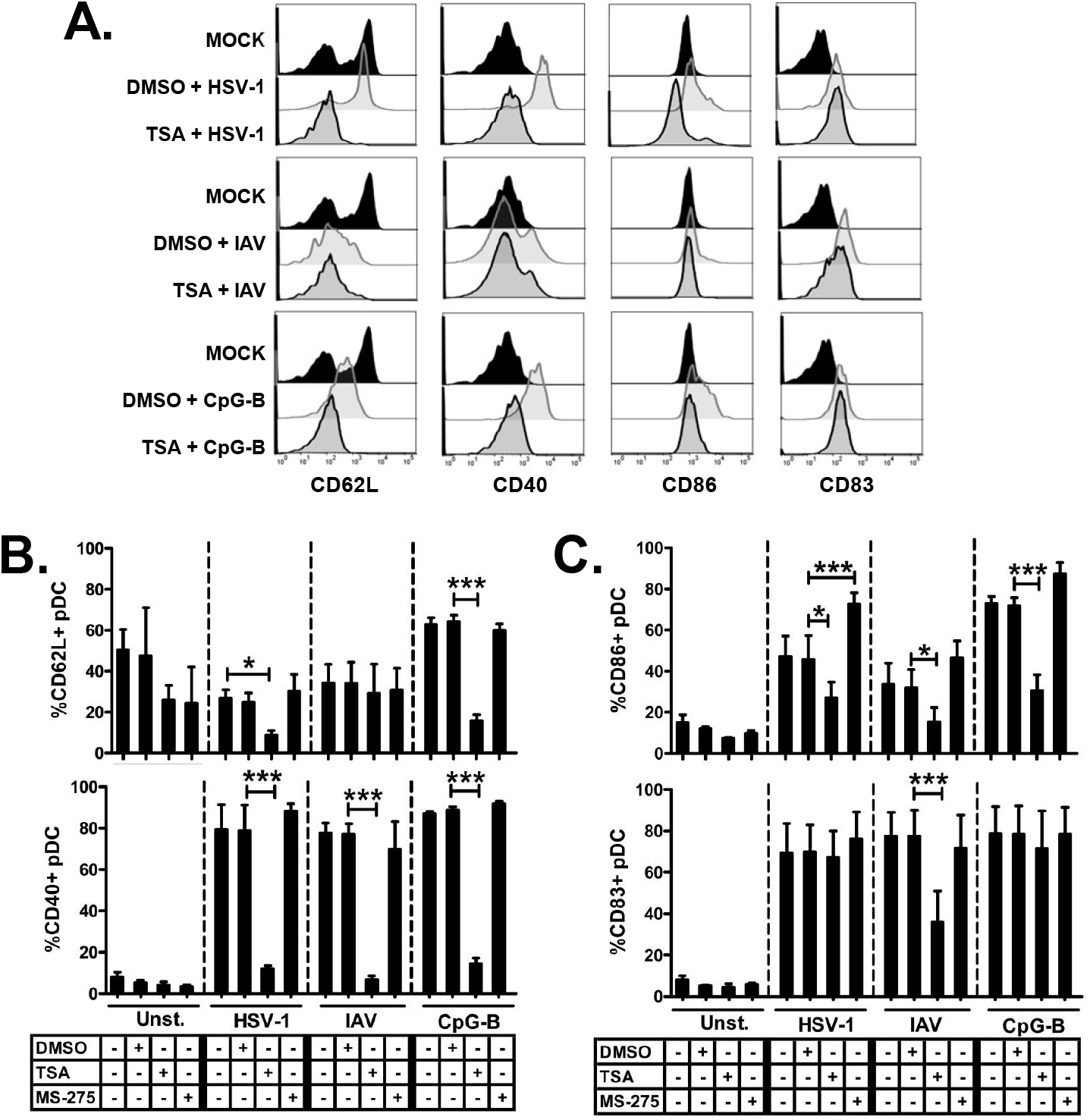
TSA and MS-275 inhibit the upregulation of pDC maturation markers upon TLR7/9 stimulation, and increase the shedding of the activation marker CD62L. PBMC were incubated with TSA (100 ng/ml), MS-275 (5 μM), or DMSO (1:530) for 1 hour and then stimulated with HSV-1 (MOI of 1), IAV (MOI of 2), or CpG-B (5 μg/ml) for 8 hours. Representative data derived from events first gated on PBMC, then pDC were identified by a BDCA-2^+^/CD123^high^ gate. From this gate, histograms were derived and gates applied based on the unstimulated control in DMSO (Mock) **(A)**. Surface expression of CD62L **(B, top)**, CD40 **(B, bottom)**, CD86 **(C, top),** CD82 **(C, bottom)** were measured by flow cytometry and reported as percentage. (N = 4, 4 independent experiments with different donors; Data are expressed as mean ± SEM; Data were analyzed with 1-way ANOVA and Bonferroni’s post test; *p<0.05; **p<0.01; ***p<0.001).

At this point, we decided to add a more clinically relevant HDACi as part of our study. SAHA, also known as Vorinostat, has been implicated in the downregulation of innate immune function in various animal models [17, 18, 25]. In addition, SAHA, marketed as Zolinza is FDA-approved for the treatment of cutaneous T cell lymphoma, and it is being utilized in over 64 active clinical trials alone or in combination with other therapies for the treatment of several cancer indications including solid tumors and hematological cancers. After following the same experimentation modality as with TSA and MS-275, we observed that SAHA also inhibited the production of IFN-α and TNF-α cytokines in a dose-dependent manner and this inhibition was statistically significant at 500 nM **(Fig. 4A-C)** without causing pDC apoptosis **(data not shown).**

**FIGURE 4.**
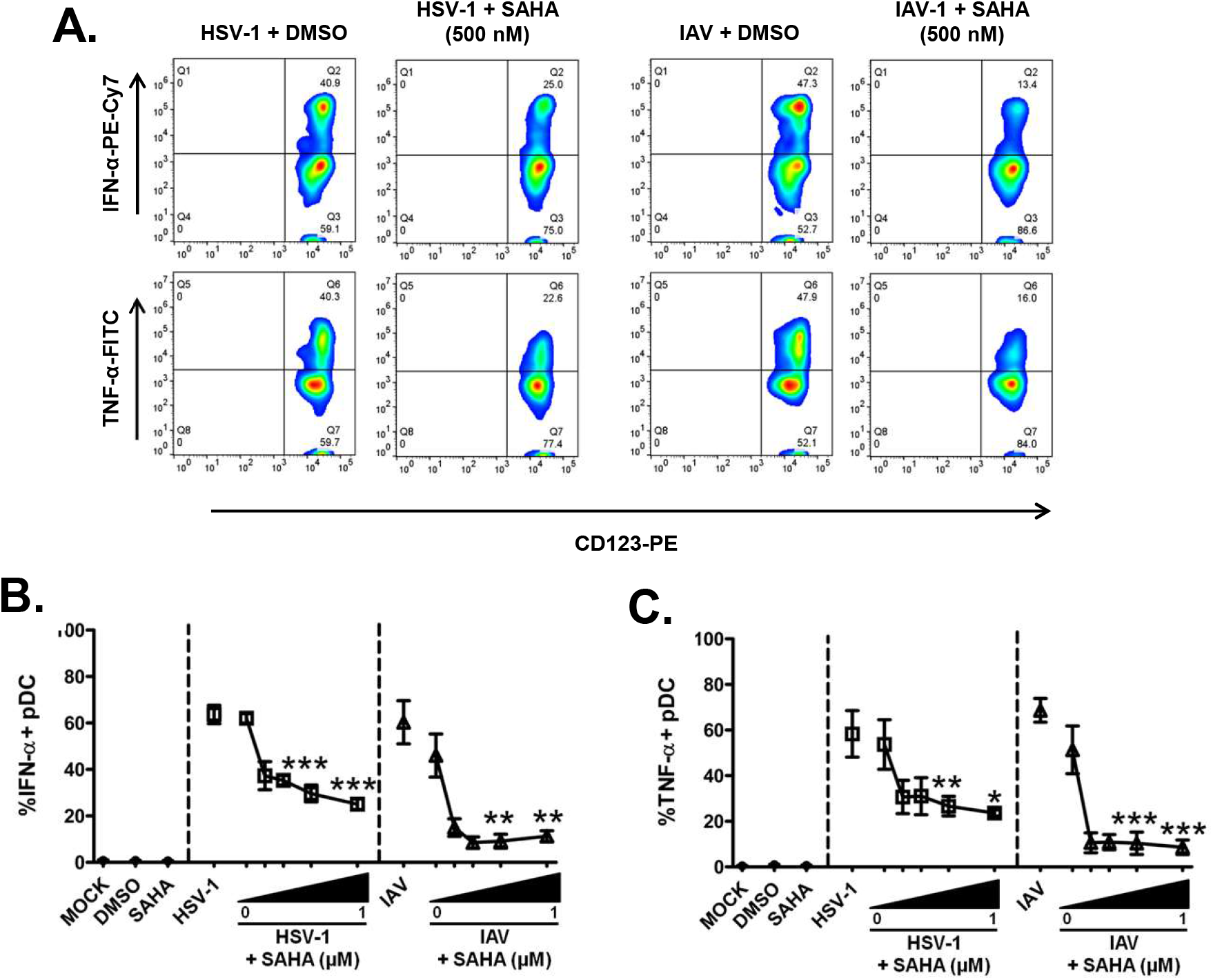
SAHA inhibits HSV-1 and IAV-induced IFN-α and TNF-α production in human pDC. PBMC were incubated with SAHA at decreasing concentrations for 1 hour and then stimulated with HSV-1 or IAV for 6 hours. Cells were then processed with intracellular flow cytometry to detect IFN-α and TNF-α production in pDC. Representative histograms showing the differential inhibition of intracellular HSV-1 (MOI of 1) and IAV-induced IFN-α and TNF-α production in pDC by SAHA (500 nM) or DMSO vehicle control (1:1000) **(A)**. Pooled data of dose curve experiments showing a dosedependent effect on HSV-1- (MOI of 1; squares) or IAV (MOI of 2; triangles) -induced IFN-α **(B)** and TNF-α **(C)** by SAHA (0-1000 μM) when compared to the respective vehicle control. (N = 3; 3 independent experiments from different donors; Data are expressed as mean ± SEM; Data were analyzed with 1-way ANOVA with Bonferroni’s post test; *p<0.05; **p<0.01; ***p<0.001).

We concluded that in most instances, treatment with a broad and a Class I HDACi can effectively inhibit TLR7/9-induced cytokine production and maturation. However, using a class I HDACi has less impact on TLR-induced pDC maturation. Moreover, the outcome can vary depending on whether TLR7 or TLR9 stimulus is used suggesting a differential effect on upstream or downstream process during signaling

### TSA and MS-275, but not SAHA, downregulate HSV-1-mediated IFN-α production without affecting IRF-7 upregulation

Plasmacytoid DC express high constitutive levels of IRF-7, which is crucial for the rapid response of human pDC to host infection and IRF-7 basal levels, are rapidly upregulated upon viral stimulation through the TLR7/9 pathways [4, 5]. Interestingly, IRF-7 expression has been shown be negatively affected by HDACi treatment in murine bone marrow-derived macrophages stimulated with TLR2/4 ligands [25]. Thus, we questioned whether the previously observed inhibition of IFN-α in the presence of TSA, MS-275, or SAHA could be associated with a detrimental effect on IRF-7 expression. Surprisingly, pre-treatment of PBMC with TSA or MS-275 had no inhibitory effect on basal levels of IRF-7 in pDC or on HSV-1- and IAV-induced upregulation of IRF-7, but consistently inhibited IFN-α production **(Fig. 5B, C)**. In contrast, SAHA inhibited IRF-7 upregulation only when IAV was utilized as a TLR agonist, but not when HSV-1 was used **(Fig. 5D).**

**FIGURE 5.**
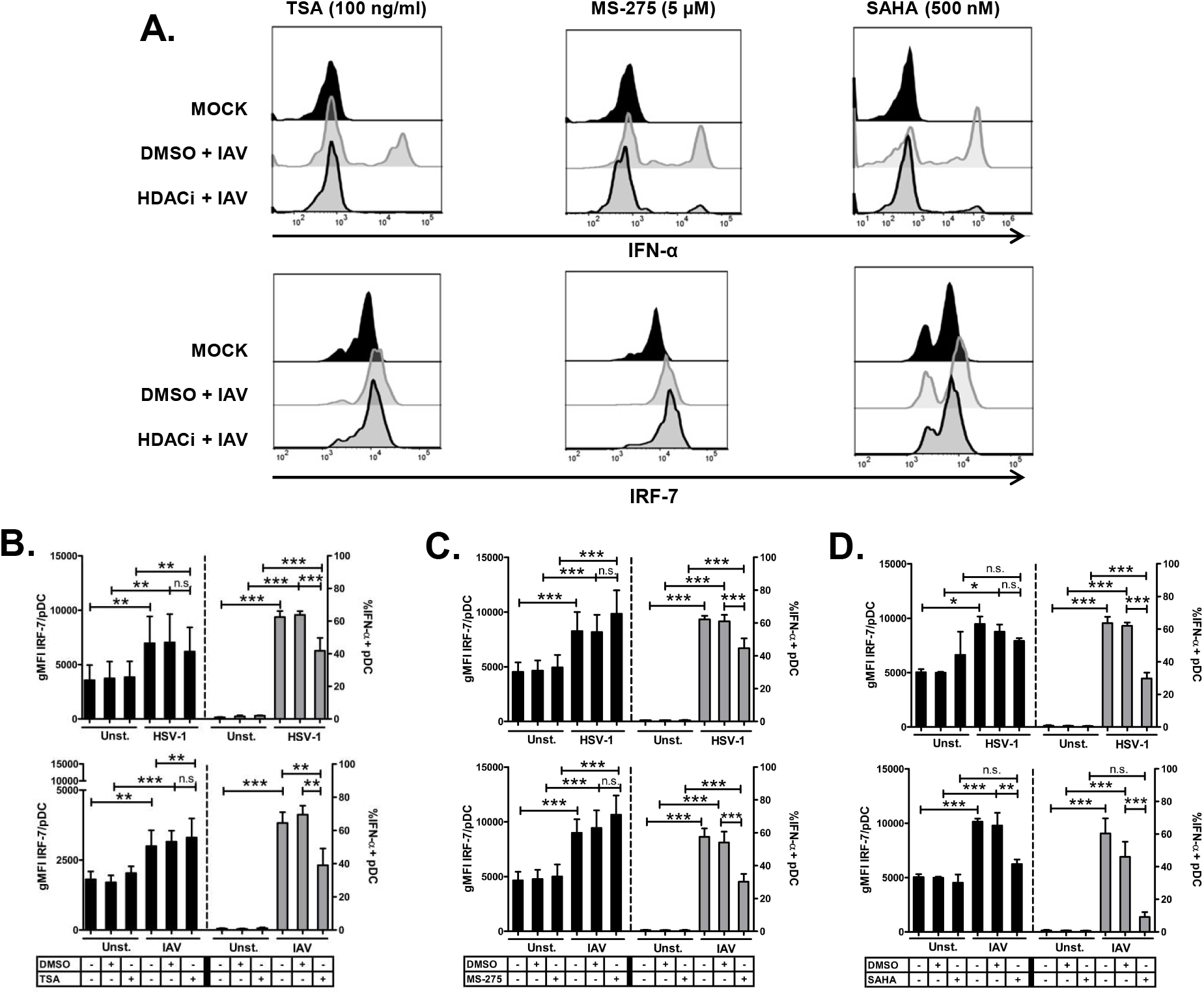
TSA and MS-275, but not SAHA, inhibits IAV and HSV-1-mediated IFN-α protein upregulation without decreasing IRF-7 upregulation. PBMC were pre-treated with TSA (100 ng/ml), SAHA (500 nM) and DMSO vehicle control (1:1000) or MS-275 (5 μM) and DMSO (1:530), for 1 hour and then stimulated with HSV-1 (MOI of 1) or IAV (MOI of 2) for 6 hours. IRF-7 (black bars) and intracellular IFN-α (gray bars) protein expression was measured concurrently by intracellular flow cytometry. Representative data showing the effect, on IAV-induced IFN **(A, top)** and IRF-7 **(A, bottom)** upregulation in pDC, by TSA, MS-275, and SAHA **(A).** Cumulative experiments showing the effect on HSV-1 (top) and IAV (bottom)-induced upregulation of IFN-α and IRF-7, by TSA (100 ng/ml) **(B)** and MS-275 (5 μM) **(C)** and SAHA (500 nM) **(D).** (N = 5; 5 independent experiments with different donors for IAV. N = 3; 3 independent experiments with different donors for HSV-1. Data are expressed as mean ± SEM; Data were analyzed with 1-way ANOVA with Bonferroni’s post test; (* = p<0.05; ** = p<0.01; *** = p<0.001).

Thus, the effect observed by TSA, MS-275, and SAHA on cytokine production was independent of detrimental effects on TLR9-induced IRF-7 upregulation as well as on IRF-7 basal levels and suggested a alternative inhibitory mechanism of pDC function by HDACi.

### TSA, MS-275, and SAHA detrimentally affect HSV-1-induced IRF-7 and NF-kB p65 nuclear translocation

Based on our previous results that showed the inhibition of IFN-α upregulation by all HDACi, we questioned whether the inhibitory effect was caused by a downregulation of the upstream regulatory pathway that naturally leads to the upregulation of cytokines and maturation markers in pDC. We first interrogated the effect of TSA, MS-275, and SAHA on IRF-7 nuclear translocation upon HSV stimulation. To this end, we utilized a pDC negative enrichment approach followed by imaging flow cytometry to quantitatively measure IRF-7 at 4 hours, which is critical for induction of IFN-α. A representative analysis of how nuclear translocation was measured using ImageStream, showing the gating strategy based on the vehicle control, is shown in **Figure 5A** with sample images of translocated and non-translocated events **(Fig. 6B)**. TSA significantly decreased the HSV-1 and IAV-induced IRF-7 nuclear translocation **(Fig. 6C, D).** Moreover, SAHA and MS-275 had a similar effect, but to a lesser extent than TSA. **(Suppl. Fig. 3)**.

**FIGURE 6.**
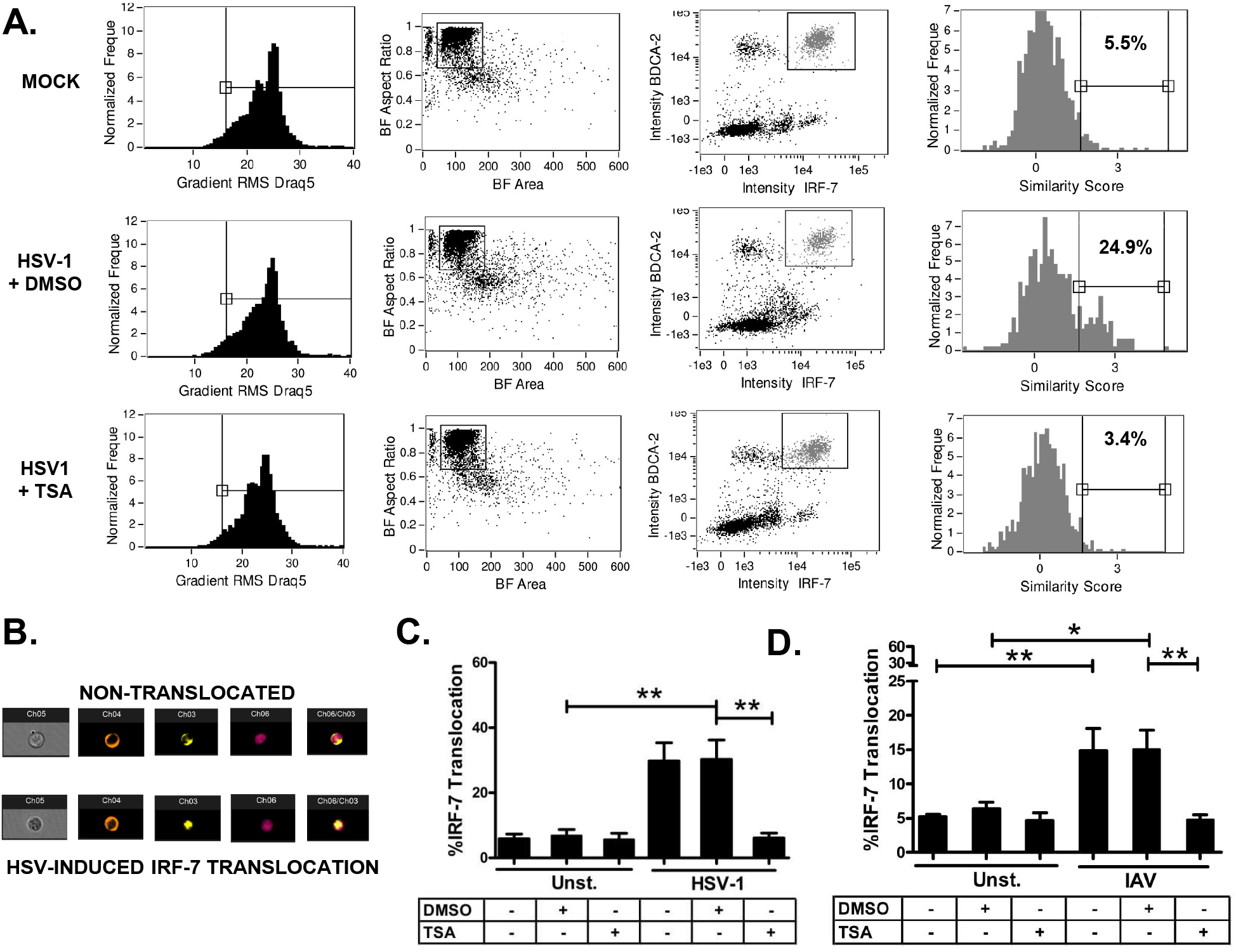
TSA decrease HSV-1 and IAV-induced IRF-7 nuclear translocation. Negatively enriched pDC were incubated with TSA (100 ng/ml = 300 nM), SAHA (1 μM), MS-275 (5 μM) or DMSO (1:1000) for 1 hour and then stimulated with HSV-1 (MOI of 1) for 4 hours. Samples were surface stained for pDC using anti-human BDCA-2 PE, fixed overnight, permeabilized, and then intracellularly stained with IRF-7 AF488 and DRAQ5 (nuclear stain), and acquired using the ImageStream flow cytometer. Single events were identified by plotting the Bright Field (BF) aspect ratio vs. area, then events in focus were gated in a gradient RMS histogram of the DRAQ5 nuclear stain, IRF-7^high^/BDCA-2^+^ pDC were gated by plotting a dot plot of the BDAC-2 vs. IRF-7 intensities, finally the percent of IRF-7 translocation was gated in a IRF-7/DRAQ5 similarity score histogram based on the DMSO only control (Mock). Representative analysis for the TSA-mediated effect on HSV-1-mediated IRF-7 nuclear translocation in pDC **(A)**. Sample images of non-translocated and translocated events are shown. BF images (Ch05), BDCA-2 PE (Ch04), IRF-7 AF488 (Ch03), DRAQ5 nuclear stain (Ch06), IRF-7/DRAQ5 merged images (Ch06/Ch03) **(B)**. Experiments showing the HSV-1 **(C)** or IAV **(D)**-induced IRF-7 nuclear translocation in the presence of TSA gates were drawn based on DMSO vehicle control. (N = 4; 4 independent experiment with different donors for TSA; Data are expressed as %mean ± SEM; Data were analyzed with 1-way ANOVA with Bonferroni’s post test; * = p<0.05; ** = p<0.01; *\*\*\** = p<0.001).

In order to gain more insight into the effects of HDACi on TLR signaling and considering the inhibitory effects on maturation markers and TNF-α production observed previously, we also interrogated the status of the NF-κB transcription factor. Following a similar treatment time and assay setup as with IRF-7; we interrogated the nuclear translocation of NF-κB p65 subunit post IκBα degradation. Representative histograms from Image Stream analysis showing the inhibitory effect on NF-κB p65 nuclear translocation in the presence of TSA and SAHA are shown in **Figure 7A**. We observed that both TSA and SAHA significantly decreased NF-κB p65 nuclear translocation **(Fig. 7B).** Interestingly, we also observed a decrease in overall NF-κB fluorescence in the presence of SAHA, but not TSA, suggesting that SAHA can affect overall NF-κB p65 protein expression in pDC **(Fig. 7C).**

**FIGURE 7.**
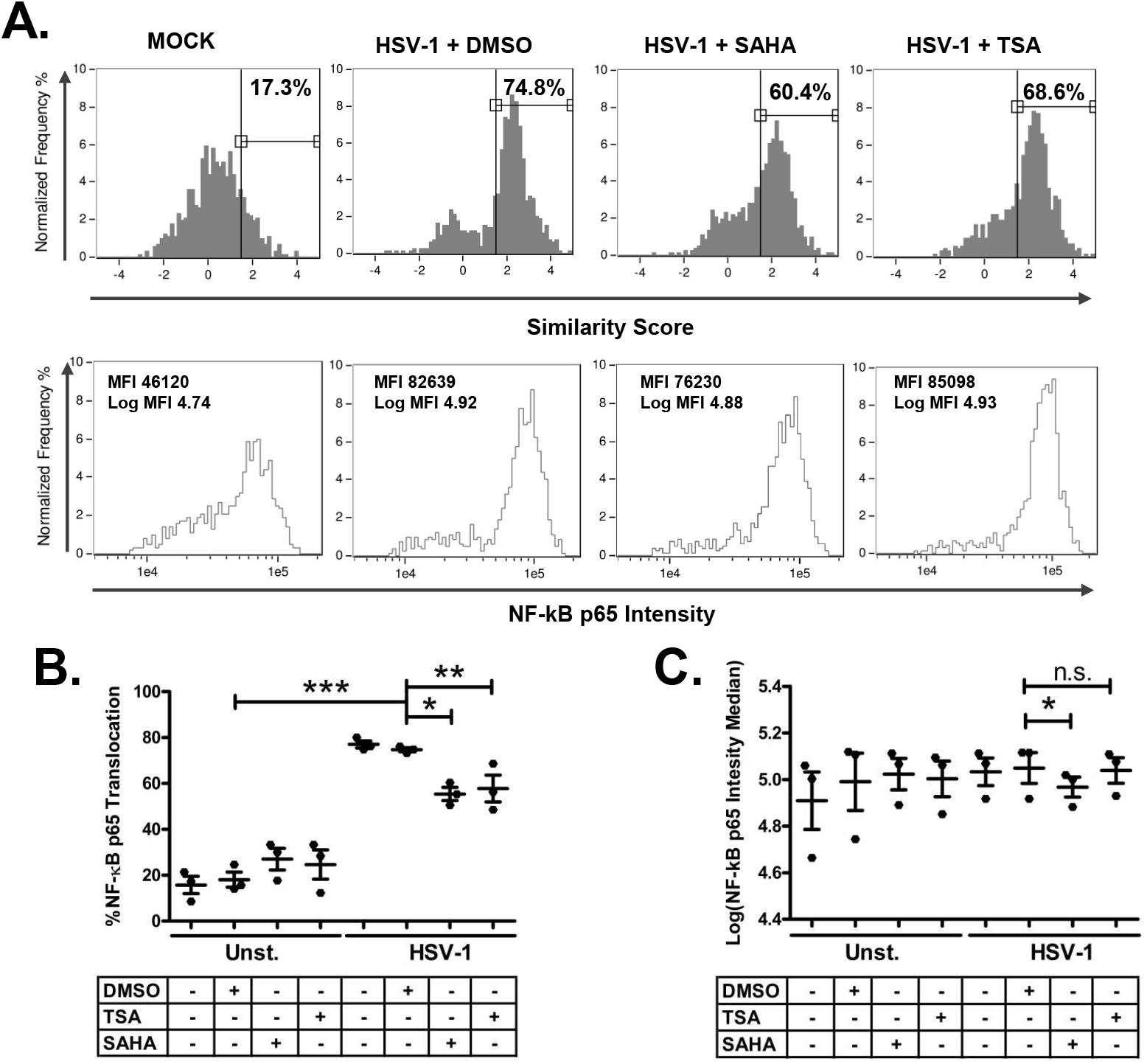
SAHA and TSA decrease TLR9-induced NF-kB p65 nuclear translocation. Representative analysis for the TSA- and SAHA-mediated effect on HSV-1-mediated IRF-7 nuclear translocation and NF-kB p65 protein upregulation in pDC. Negatively enriched pDC were incubated with TSA (100 ng/ml = 300 nM) and SAHA (500 nM) or DMSO (1:1000) for 1 hour and then stimulated with HSV-1 (MOI of 1) for 4 hours. Samples were surface stained for pDC using anti-human BDCA-2 PE and then intracellularly stained with IRF-7 AF488 and DRAQ5 (nuclear stain), and acquired using the ImageStream and analyzed with IDEAS software. Single events were identified by plotting the Bright Field (BF) aspect ratio vs. area, then events in focus were gated in a gradient RMS histogram of the DRAQ5 nuclear stain, NF-kB p65^+^/BDCA-2^+^ pDC were gated by plotting a dot plot of the BDAC-2 vs. NF-kB p65 intensities, finally the percent of NF-kB p65 translocation was gated in a NF-kB p65/DRAQ5 similarity score histogram **(A, top).** Concurrently, NF-kB p65 overall intensity was measured from the NF-kB p65^+^/BDCA-2^+^ pDC population and the MFI was reported as log(MFI) **(A, bottom)**. Pooled experiments showing the effect of TSA and SAHA in NF-κB p65 nuclear translocation **(B)** and protein expression **(C).** (N = 3; 3 independent experiments with different donors; Data are expressed as %mean ± SEM; Data were analyzed with 1-way ANOVA with Bonferroni’s post test; * = p<0.05; ** = p<0.01; *** = p<0.001).

We concluded that the previously observed TSA and SAHA-mediated inhibition of the production of IFN-α and TNF-α can be partially explained by a failure of these transcription factors to translocate into the nucleus in the presence of TSA, MS-275and SAHA. In addition, the data suggest that upstream signaling pathways are affected by HDACi in TLR-mediated pDC activation.

### HSV-1-induced IRF-7 and NF-kB p65 activation is decreased in the presence of TSA, MS-275, and SAHA

Given that the activation of IRF-7 and NF-κB is upstream of nuclear translocation, we hypothesized that TSA and SAHA can disrupt the activation of TLR-mediated signaling cascade by negatively impacting the phosphorylation of IRF-7 and NF-κB p65. In order to answer this question, PBMC were isolated and pre-treated with TSA or SAHA for 1 hour, then stimulated with HSV-1 for 3 hours, then we utilized BD Phosflow^™^ assays to assess the phosphorylation status of IRF-7 and NF-κB p65 in the presence of TSA, MS-275, SAHA.

An example of the BD Phosflow^™^ cytometry analysis **(Suppl. Fig. 4)** shows our switching from anti-human BDCA-2 to HLA-DR antibodies since the BDA-2 antibody is sensitive to the fixation method required for Phosflow^™^

Using HLA-DR and CD123 mAbs for detection of pDC, as well as a mAb that recognizes phosphorylated serines 477 and 479 of IRF-7 and a polyclonal antibody that recognizes the phosphorylated p65 subunit of NF-κB, a representative analysis using IAV as a TLR agonist shows the effect of HDACi on these transcription factors is shown in **Fig. 8A**. We determined that TSA, but not MS-275 or SAHA, partially reduced the phosphorylation of these transcription factors when HSV-1, but not IAV, was the stimulus **(Fig. 8B-D, top panels)**. In contrast, when IAV was the stimulus, IRF-7 and NF-κB p65 phosphorylation was markedly decreased in the presence of all HDACi **(Fig. 8B-D, bottom panels).**

**FIGURE 8.**
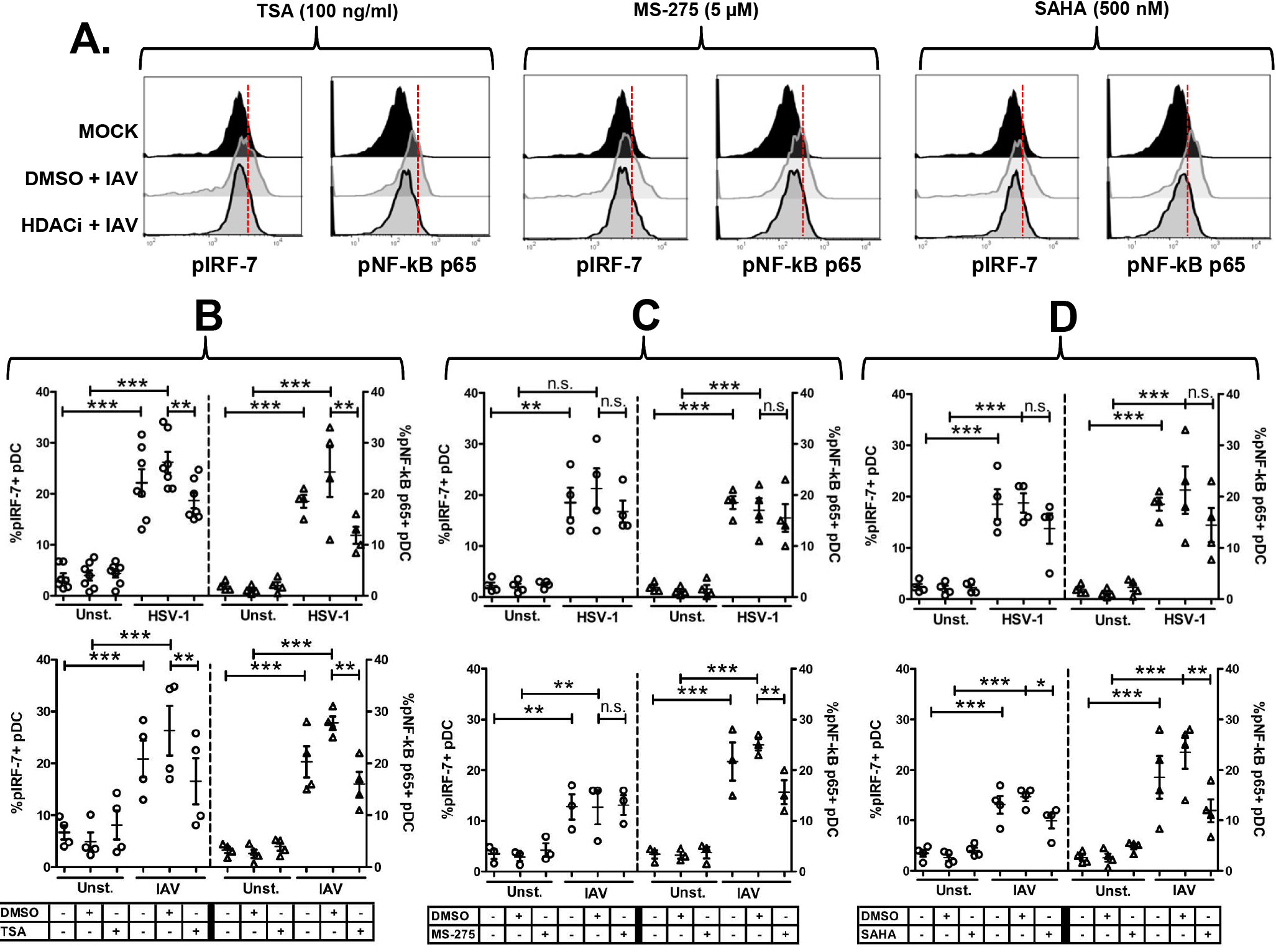
TSA, SAHA, and MS-275 decrease TLR9-induced activation of IRF-7 and NF-κB p65 transcription factors. PBMC were incubated with TSA (100 ng/ml) or DMSO (1:1000) for 1 hour, stimulated with HSV-1 (MOI of 1) or IAV (MOI of 2) for 3 hours. Samples were processed using BD Phosflow protocol. PBMC were gated based on SSC vs. FSC-A (Area) and pDC were identified by gating on +HLA-DRhigh/+CD123high events on a HLA-DR APC vs. CD123 PE intensity dot plot, then percent phosphorylated IRF-7+ pDC were gated based on mock sample for HSV-1 or IAV from which percentages were reported. Representative analysis for TSA, SAHA, and MS-275-mediated effect on IAV-mediated IRF-7 and NF-kB p65 phosphorylation in pDC. Red lines represent the position where the gate was placed from which percentages were derived. **(A)**. Pooled data with HSV-1 stimulation (top panels) and IAV (bottom panels) showing the effect of TSA (100 ng/ml) **(B)**, MS-275 (5 μM) **(C)**, and SAHA (500 nM) **(D)**, or DMSO (1:1000) on the phosphorylation of IRF-7 and NF-kB p65. Data represent the percentage of pIRF-7+ and pNF-κB p65+ pDC with gating based on mock sample. (N = 4-5, 4-5 independent experiments with different donors; Data are expressed as mean ± SEM; Data were analyzed with 1-way ANOVA with Bonferroni’s post test; n.s. = not significant).

Thus, the effect of HDACi on transcription factor activation can vary depending on the TLR pathways, 7 or 9, that is activated in human pDC. These and our previous results show a selectivity for signaling pathways affected by HDACi.

## Discussion

Our current study explored the effects of TSA and SAHA, two broad class I and II HDACi, and MS-275, a class I HDACi on human pDC biology. We observed that all HDACi used in this study exerted a deleterious effect on IFN-α and TNF-α production upon viral TLR7/9 stimulation. In addition, TLR-mediated upregulation of maturation markers was observed for TSA, but not MS-275. These effects were strongly associated with disturbances in IRF-7 and NF-kB transcription factor nuclear translocation and phosphorylation.

We were particularly interested on the effect by HDACi during early pDC stimulation because pDC are the primary IFN producers during viral infection and any treatment that affects pDC would inevitably have a direct repercussion on the anti-viral first line defenses. Our 6-hour intracellular experiments revealed that TSA and MS-275 all inhibited HSV-1 and IAV-mediated IRF-7 upregulation without decreasing IRF-7 basal levels, but this was not the case for SAHA which was associated with a decline in said transcription factor upregulation. The effects observed for SAHA, but not TSA or MS-275, in IFN production could partially be explained by a decline in the upregulation of IRF-7. However, this decline could not have been due to a decrease in IRF-7 mRNA given that basal protein levels were stable. Interestingly, this scenario only affected IAV, but not HSV-1,-based stimulation indicating that TLR7 signaling is more susceptible to SAHA treatment than TLR9 signaling.

Previously, it was shown that TSA negatively affected the IFN-stimulated upregulation of IRF-7 expression by inhibiting the formation of the interferon-stimulated gene 3 (ISGF3) complex (composed of STAT1/STAT2/IRF-9 proteins) and this was associated with impairment of STAT2 nuclear accumulation, but not STAT1, in mouse L929 cells [26]. Likewise, we previously reported the same effect by TSA in human pDC when using recombinant IFN-α to trigger the IFN receptor (IFNR) [27]. Hence, in the current study the observed inhibition of total IFN-α production at 24 hours by TSA could partially be attributed to the interruption of signaling events in the IFNR axis; while MS-275 remains to be investigated in this regard. On the other hand, stimulation of TLR7 or 9 in the presence of TSA and MS-275 did not inhibit upregulation of IRF-7, yet it blocked cytokine production. We can infer from this finding that HDACi effects on transcription factor expression may not be the only inhibitory pathway that leads to inhibition of pDC biology.

TSA, but not MS-275 exhibited dramatic effects on TLR7/9-induced pDC upregulation to maturation markers and activation. We did not expect to observe a complete lack of inhibition, and in some instances, an increase in maturation marker expression by MS-275. Nencioni *et al*. [24] showed that nanomolar concentrations of MS-275 inhibited poly I: C-induced CD86/CD83 upregulation of IL-4/GM-derived human MDDC in a dose-dependent manner. Even though we used MS-275 at a concentration 125 times higher than the highest dose used by Nencioni *et al*. for MDDC, it was not strong enough to negatively affect pDC maturation. The concentration used in our studies was based on studies by Nebbioso *et al*. [28], which showed that MS-275 at 5 μM specifically affected canonical class I HDACs activity in an *in-vitro* system for testing of HDAC enzymatic activity [28]. Hence, we propose that in human pDC the regulation of Class II HDACs is important for TLR7/9 signaling events that are critical for pDC maturation possibly via the MAP-kinase pathway. Moreover, the observed effect could also be attributed to HDACi potency, which has been shown to be stronger for TSA and SAHA than for MS-275 as evidence by the acetylation of tubulin in MCF-7 cells [29].

Interestingly, the inhibition of IAV-induced CD83 upregulation was not observed for HSV-1 and CpG-B-induced CD83 upregulation by TSA, which suggests that HDACi effect, may be dependent on the nature of the TLR agonist. It remains to be explored whether a TLR7 ligand, other than IAV, such as synthetic 3M-003, which has been shown in our laboratory to induce IFN-α without inducing IRF-7 translocation (unpublished results), would result in a similar inhibitory profile with TSA or SAHA. While exploring the effects on the pDC homing receptor ligand CD62L, we discovered a marked decreased on this ligand on the pDC surface in the presence of TSA. The absence of CD62L on the pDC surface could either mean a decrease protein output or an accelerated state of shedding. In either case, the effect would result in incapacity by pDC to home to secondary lymphoid tissues and elicit an immune response during a viral infection.

Because of the observed TSA-induced effects on IFN-early post stimulation at 6 hours and later on activation and maturation markers, we questioned the effects of TSA and SAHA on IRF-7 and NF-κB nuclear translocation. IRF-7 nuclear translocation is the hallmark signaling event that leads to IFN-α production in pDC [3]. Moreover, nuclear translocation of NF-κB is known to induce the expression of TNF-α and pro-inflammatory cytokine production [30]. We observed a very strong inhibition of IRF-7 nuclear translocation, and a moderate inhibition of NF-κB p65 nuclear translocation, at 4 hours of HSV-1 or IAV stimulation in the presence of TSA or SAHA. Our results are consistent with those reported by Salvi *et al*. [20], which showed a mechanism of inhibition by evidence of decreased IFN-α mRNA levels and IRF-7 nuclear translocation upon synthetic bacterial TLR-9 agonist using a qualitative approach [20]. We strongly believe our study formulates a clearer picture of a possible mechanism of inhibition by the following criteria: first, we used pDC that were not cross-linked via the BDCA-4 receptor by positive enrichment which can impact pDC function; second we used a more biologically relevant viral model of stimulation not shown before in combination with TSA, SAHA, and MS-275 in human pDC; third, weuse a clinically relevant HDACi, SAHA; fourth the dissection of additional IRF-7 and NF-κB signaling effects by their phosphorylation and nuclear translocation changes in the presence of HDACi. The dramatic TSA-mediated inhibitory effect on virus-induced IRF-7 translocation did not correlate with the incomplete inhibition of IFN-α production. We reasoned that TSA may affect other transcription factors, such as IRF-5, which has also been shown to be involved in the induction of IFN-α gene expression. Interestingly, IRF-5 is activated via phosphorylation by RIP2, a kinase known to function downstream of NOD2 signaling [31] and IRF-5 phosphorylation was associated with CBP recruitment and subsequent acetylation of IFNA promoter, which was blocked by TSA [32]. Thus, it is possible that inhibition of IRF-5 activity by TSA could be responsible for the unaccounted inhibition of TLR7/9-induced cytokine production, but further experimentation is required to answer this question.

Not only was type I IFN strongly inhibited by TSA and SAHA, but also TNF-α; a pro-inflammatory cytokine, which acts on other immune cells and different tissues, like the liver and hypothalamus, inducing fever and an acute phase responses. While we have shown here that TSA and SAHA-mediated inhibition of NF-κB p65 activation is indeed one of the causes for TNF-α downregulation, it is possible that the inhibitory effect on TNF-α production may also be caused by a direct effect on the regulation of NF-κB mRNA transcript production. Chen *et al*.[33] showed that mutations to S536 and S276 in NF-κB p65 (RelA) negatively affected acetylation of K310, and this in turn negatively affected RelA expression [33]. In addition, SAHA has been shown to decrease NF-κB/DNA binding in cancer cell lines, which was associated with a decrease in the production of TNF-α protein [34]. Finally, we also observed a moderate and statistically significant reduction in NF-kB p65 protein expression by ImageStream flow cytometry suggesting a possible reduction in the NF-kB p65 mRNA levels.

Recently, van Noort *et al*. elegantly and clearly showed that a cross-talk between acetylation and phosphorylation exists in prokaryotes [35]. Their findings prompted us to ask the question whether TSA, MS-275, or SAHA negatively affect the activation state of IRF-7 and NF-κB in primary pDC. Our results showed that TSA moderately, and partially, affected total amount of phosphorylated IRF-7 in pDC. In our studies, the phosphorylation of IRF-7 by BD Phosphoflow never reached 100% of pDC at 3 hours of stimulation with IAV and HSV-1, neither did a shorter (1 hour) or a longer (5 hour) time point increase the phosphorylation level IRF-7 higher than our current observations at 3 hours (data not shown); suggesting that the induction of IFN-α is not solely due to the activation of IRF-7, but possibly due to other post translational modifications on IRF-7. Interestingly, nuclear acetylation of IRF-7 has been demonstrated to occur by Caillaud *et al*. in another cell system and its acetylation promoted DNA binding to the ISG [36]. Given that acetylation and deacetylation of proteins are widespread processes that also extend to the cytoplasm; we propose that a HDACi can affect the acetylation/phosphorylation balance causing a deregulation of IRF-7 and NF-κB activation.

It is clear from our IRF-7 and NF-kB activation study that the nature of the TLR agonist plays a role. For example, TSA indiscriminately inhibited both HSV-1 and IAV action of said transcription factors, SAHA inhibition when IAV was present was statistically significant, and shows a trend when HSV-1 was used, which was not statistically significant. This difference is more obvious with MS-275 for which it is clear that its strongest and clearest pattern were observed when IAV was used and only for NF-kB phosphorylation, but not IRF-7. Yet, MS-275 inhibited IRF-7 translocation indicating that the inhibition in translocation can also be caused by other unknown mechanism beside an effect on IRF-7 phosphorylation. Currently, it is unclear how TSA, SAHA, and MS-275 moderately inhibited the activation of IRF-7 and NF-κB p65 upon TLR stimulation; however, the negative impact of HDACi on pDC nibbling may offer an explanation (data not shown). PDC nibbling is a hallmark process that leads to pDC activation via TLR signaling. It is via pDC nibbling that exogenous material is endocytosed into endosomal TLR7+/9+ compartments. Since endosomal compartments are important sites of RNA-TLR7 and DNA-TLR9 interactions, and these interactions are dependent on F-actin dynamics-as they are inhibited by F-actin polymerization inhibitor Cytochalasin-D, it is possible that inhibition of the pDC nibbling process by HDACi is affecting downstream TLR7 and −9 signaling. Considering that TSA has a high affinity for HDAC6 as a target [37], and has been shown to target cortactin, a F-actin related molecule whose acetylation controls its binding to F-actin and its dynamics [13]. That explanation may suffice for TSA, however, our data shows that MS-275 (data not shown), which is known to have no effect on HDAC6 activity, but HDAC1, 2, and 3 instead, was also able to inhibit pDC nibbling suggesting a different inhibitory pathway for MS-275 that excluded uptake.

In conclusion, we have demonstrated that HDACs play a very important role in the TLR7/9-induced function in human pDC. Our data demonstrate that HDACi, TSA, MS-275, and SAHA have quite detrimental effects on pDC cytokine production, maturation, and nibbling abilities caused by partial interruption of TLR7/9 signaling in human pDC. Our work complements the growing literature that demonstrates the negative effects of HDACi on innate immunity, especially at it relates to virus-induced pDC activation. Importantly, as HDACi are making their way into the clinic to reactivate HIV in a shock and kill strategy, it may be prudent to consider the addition of a co-therapy to aid the innate arm of the immune system such as supplementation of IFN during HDAC treatment.

## Acknowledgements

We thank Drs. Elizabeth Raveche, Christine Rohowsky-Kochan, and Theresa Chang (Rutgers University Research Faculty) for their scientific discussions. We thank Ms. Katherine Lamauro for assisting in completing final manuscript revisions and corrections (Research Intern at JTCC-Hackensack-Meridian)

## Author Contributions

Conceived and designed the experiments: DBDM, JD, and PFB. Performed experiments:

DBDM. Analyzed the data: DBDM. Provided technical and analytical expertise with Imaging Flow cytometry: SS and PFB. Wrote manuscript: DBDM. Edited manuscript: PFB and JD.

