## Supplementary Figure 1 for "HDAC inhibitors decrease TLR7/9-mediated human plasmacytoid dendritic cell activation by interfering with IRF-7 and NF-κB signaling"

### Slide 1
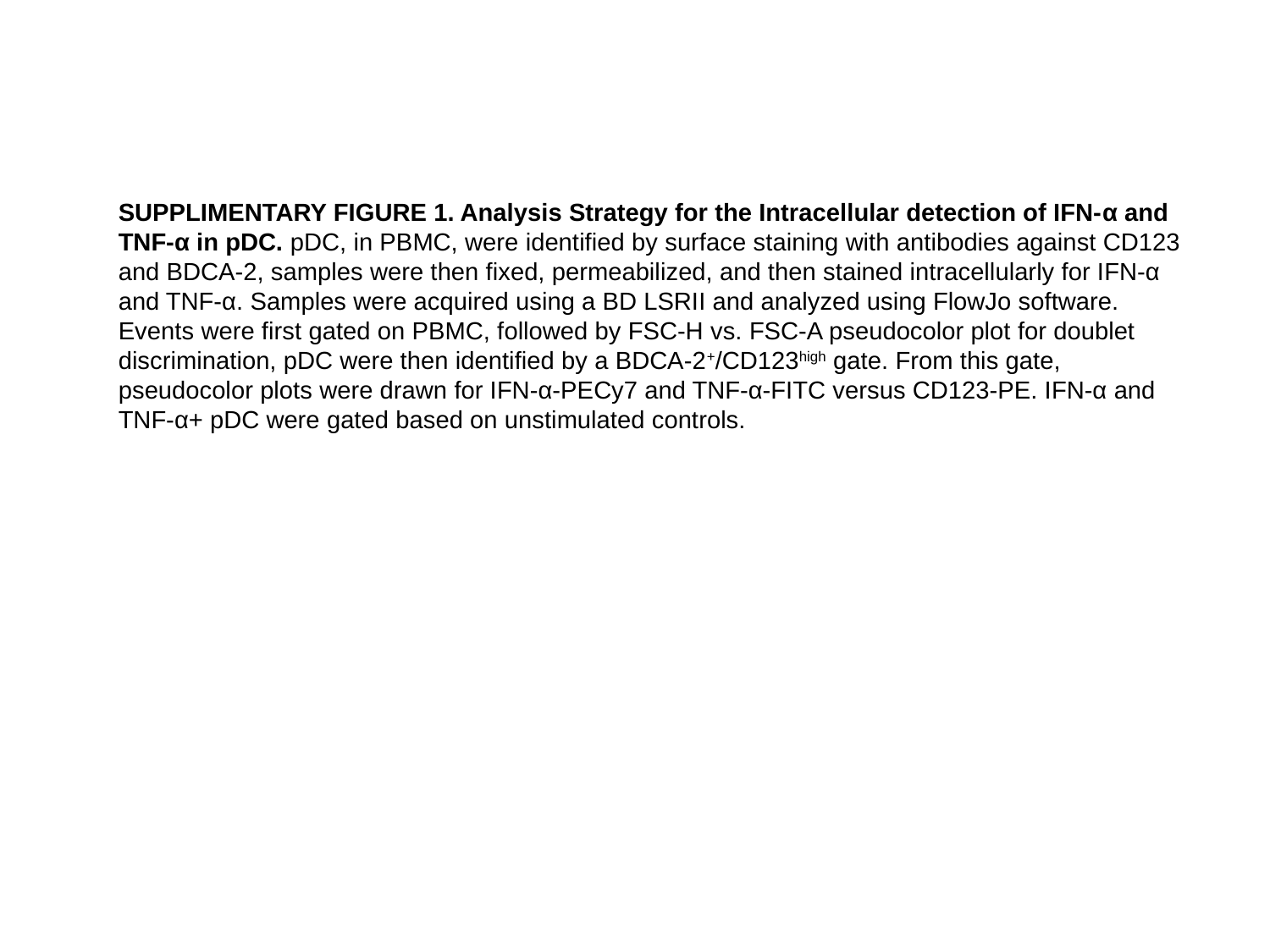

SUPPLIMENTARY FIGURE 1. Analysis Strategy for the Intracellular detection of IFN-α and TNF-α in pDC. pDC, in PBMC, were identified by surface staining with antibodies against CD123 and BDCA-2, samples were then fixed, permeabilized, and then stained intracellularly for IFN-α and TNF-α. Samples were acquired using a BD LSRII and analyzed using FlowJo software. Events were first gated on PBMC, followed by FSC-H vs. FSC-A pseudocolor plot for doublet discrimination, pDC were then identified by a BDCA-2^+^/CD123^high^ gate. From this gate, pseudocolor plots were drawn for IFN-α-PECy7 and TNF-α-FITC versus CD123-PE. IFN-α and TNF-α+ pDC were gated based on unstimulated controls.

### Slide 2
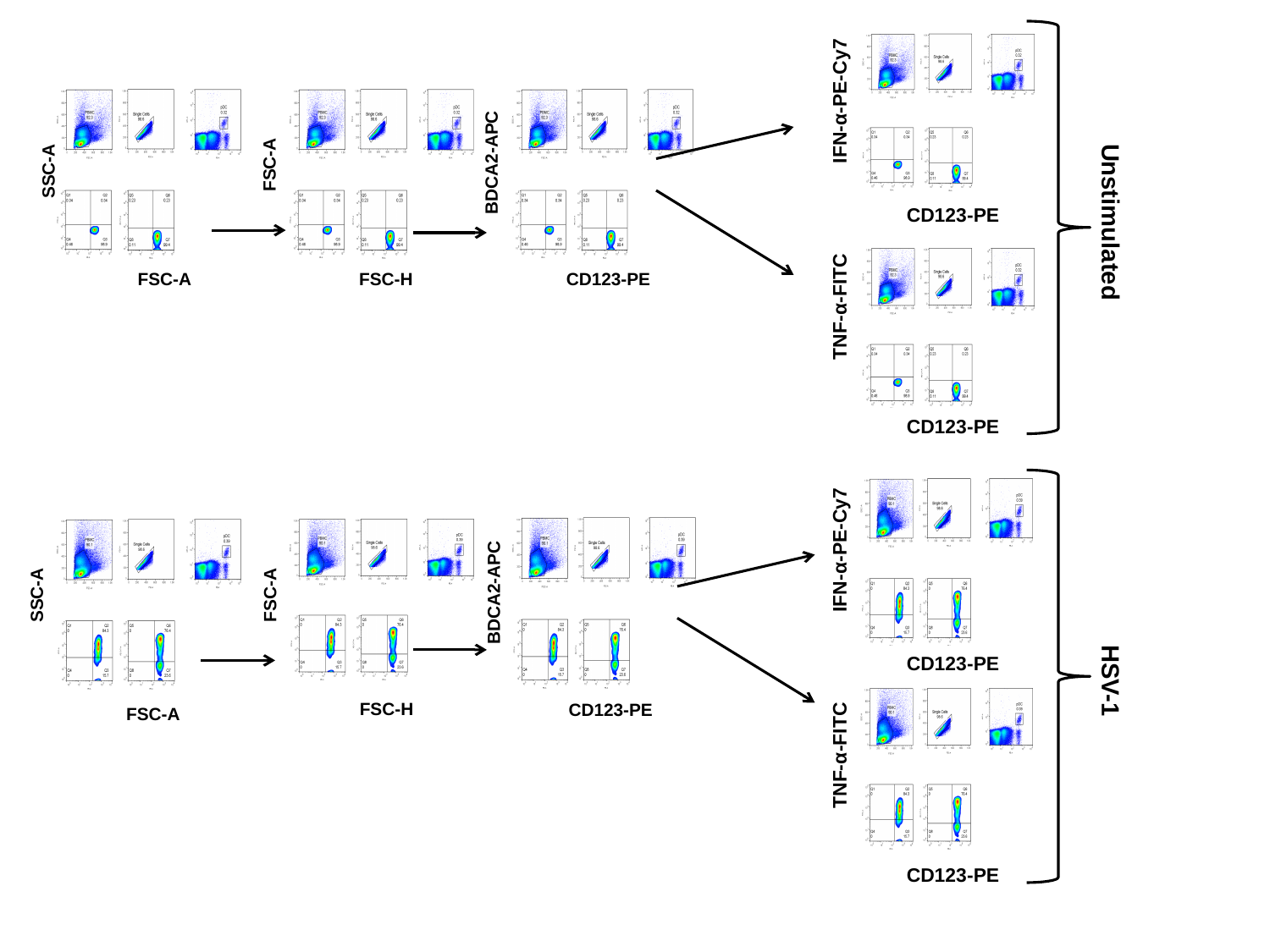

IFN-α-PE-Cy7
BDCA2-APC
FSC-A
SSC-A
CD123-PE
Unstimulated
FSC-H
FSC-A
CD123-PE
TNF-α-FITC
CD123-PE
IFN-α-PE-Cy7
BDCA2-APC
SSC-A
FSC-A
CD123-PE
HSV-1
FSC-H
CD123-PE
FSC-A
TNF-α-FITC
CD123-PE
