## Supplementary Figure 3 for "HDAC inhibitors decrease TLR7/9-mediated human plasmacytoid dendritic cell activation by interfering with IRF-7 and NF-κB signaling"

### Slide 1
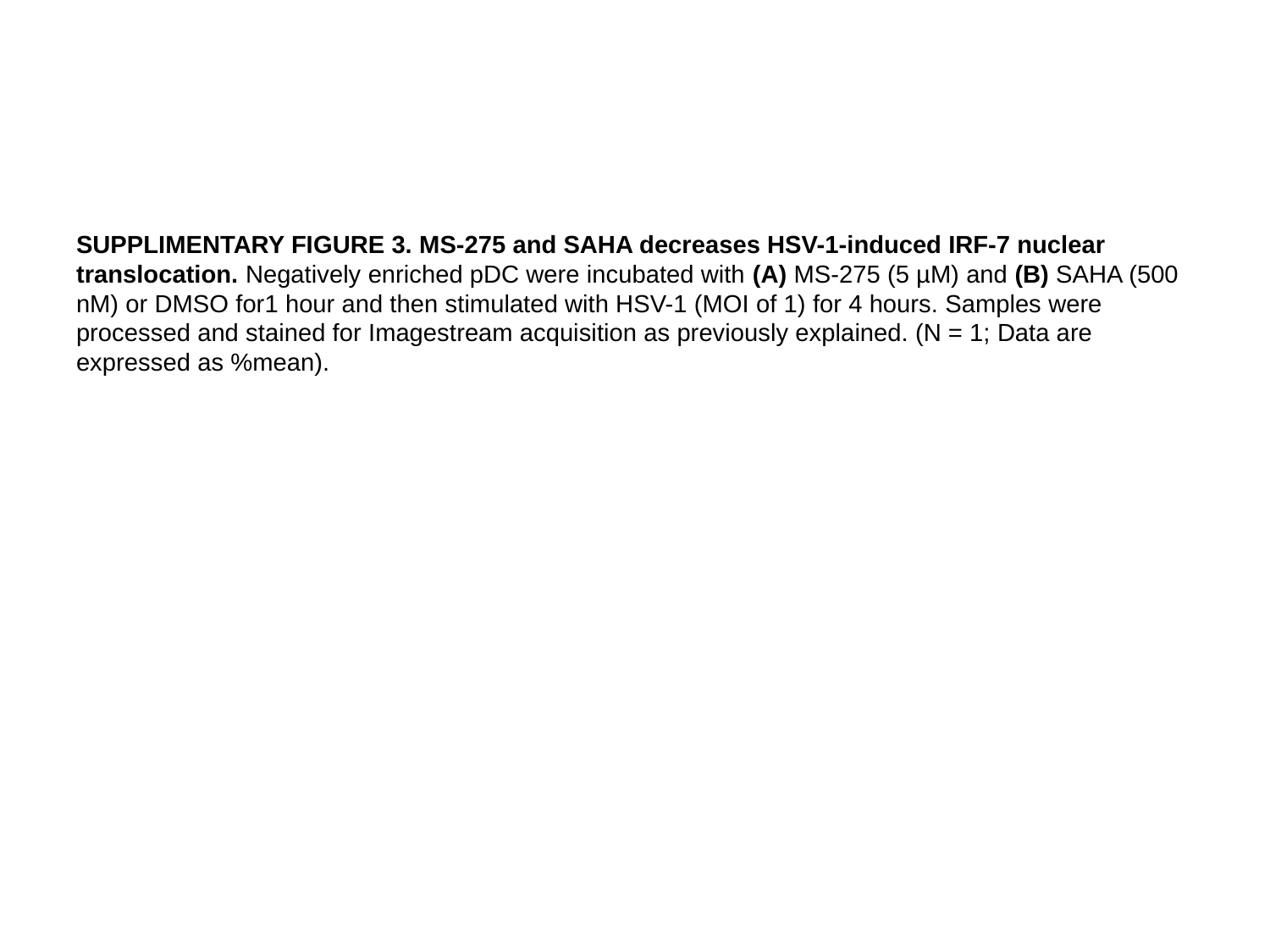

SUPPLIMENTARY FIGURE 3. MS-275 and SAHA decreases HSV-1-induced IRF-7 nuclear translocation. Negatively enriched pDC were incubated with (A) MS-275 (5 µM) and (B) SAHA (500 nM) or DMSO for1 hour and then stimulated with HSV-1 (MOI of 1) for 4 hours. Samples were processed and stained for Imagestream acquisition as previously explained. (N = 1; Data are expressed as %mean).

### Slide 2
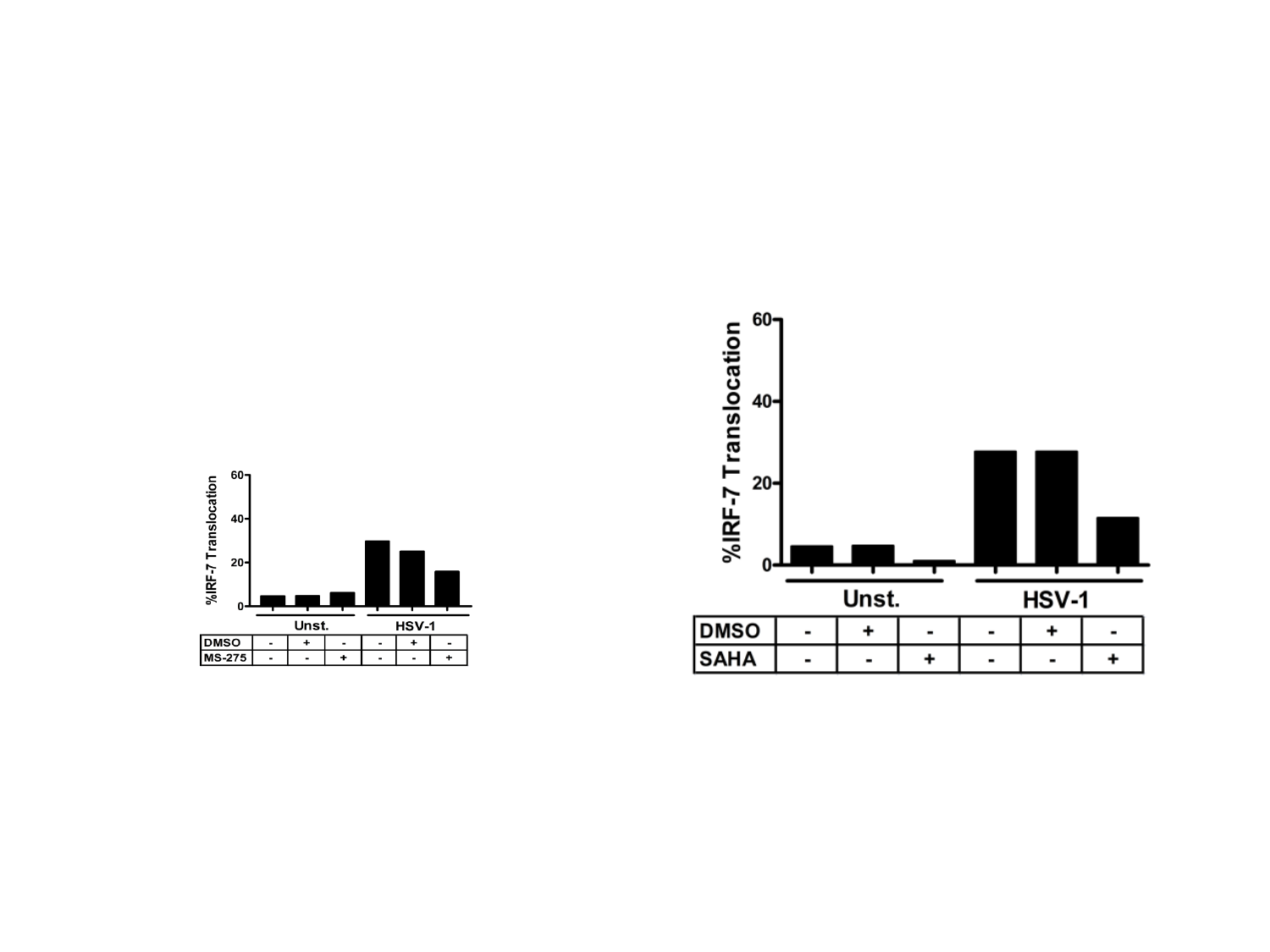
