## Supplementary Figure 4 for "HDAC inhibitors decrease TLR7/9-mediated human plasmacytoid dendritic cell activation by interfering with IRF-7 and NF-κB signaling"

### Slide 1
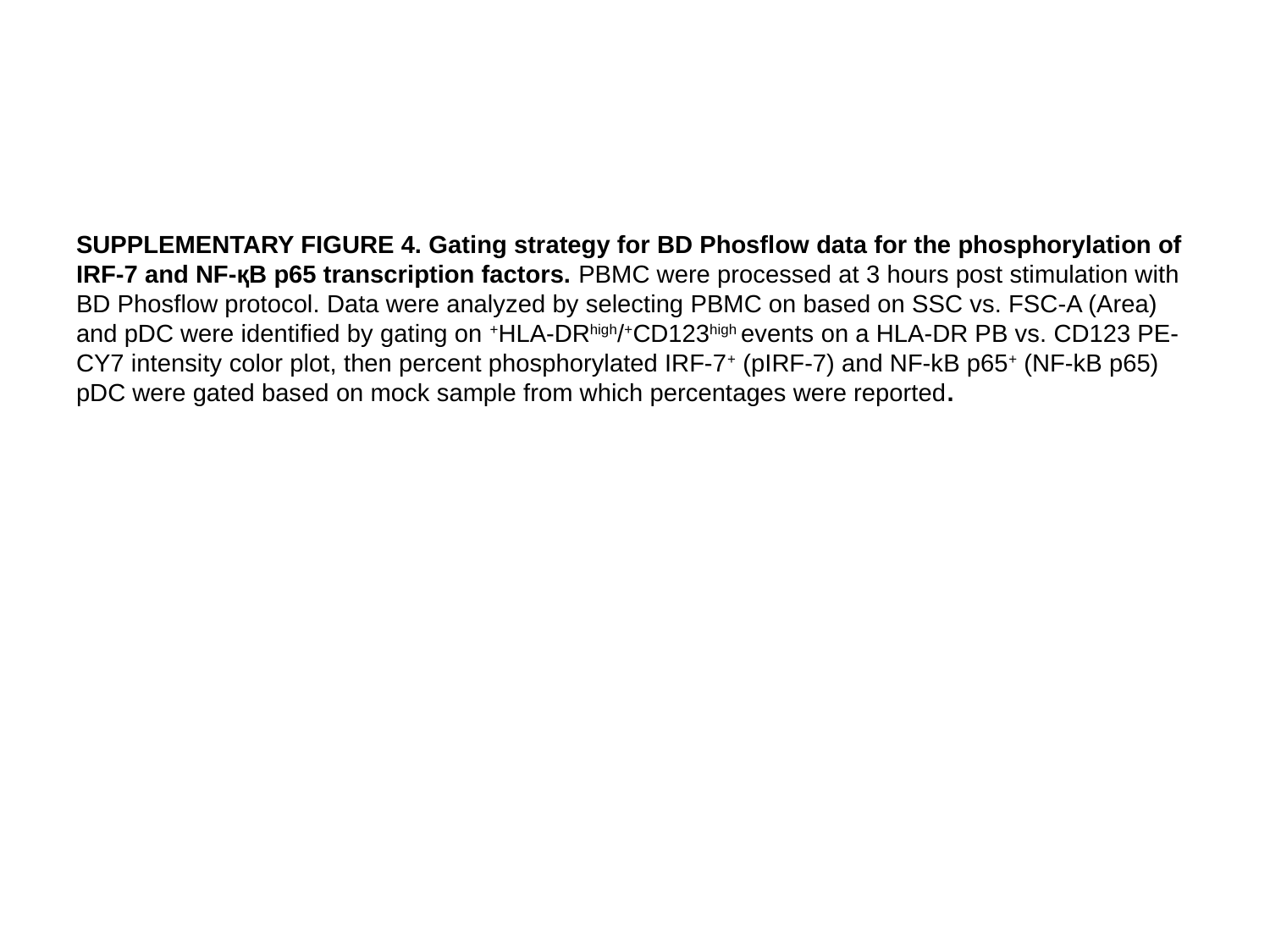

SUPPLEMENTARY FIGURE 4. Gating strategy for BD Phosflow data for the phosphorylation of IRF-7 and NF-қB p65 transcription factors. PBMC were processed at 3 hours post stimulation with BD Phosflow protocol. Data were analyzed by selecting PBMC on based on SSC vs. FSC-A (Area) and pDC were identified by gating on ^+^HLA-DR^high^/^+^CD123^high^ events on a HLA-DR PB vs. CD123 PE-CY7 intensity color plot, then percent phosphorylated IRF-7^+^ (pIRF-7) and NF-kB p65^+^ (NF-kB p65) pDC were gated based on mock sample from which percentages were reported.

### Slide 2
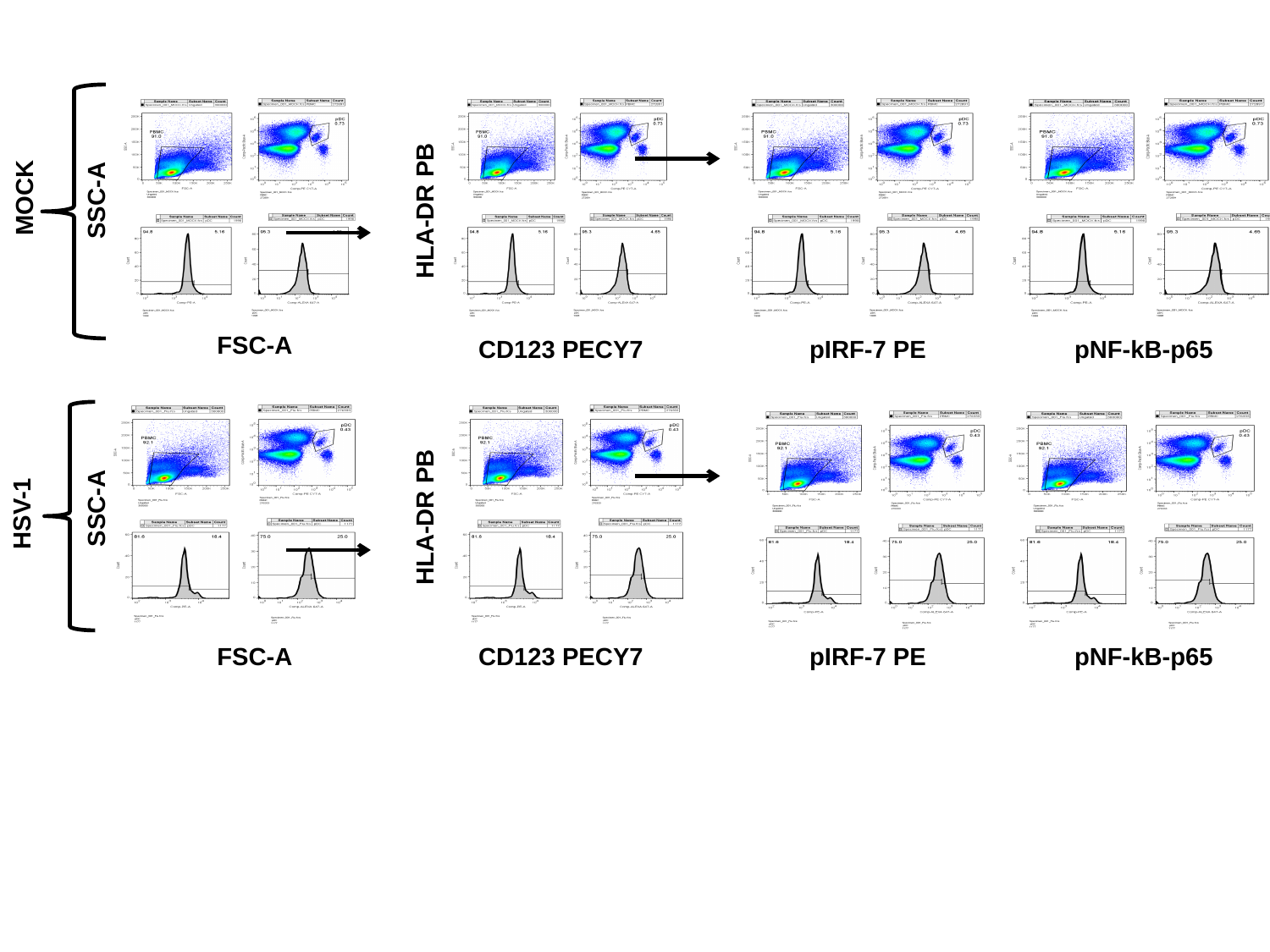

SSC-A
HLA-DR PB
FSC-A
CD123 PECY7
pIRF-7 PE
pNF-kB-p65
SSC-A
HLA-DR PB
FSC-A
CD123 PECY7
pIRF-7 PE
pNF-kB-p65
MOCK
HSV-1
