## Supplementary Figure 2 for "HDAC inhibitors decrease TLR7/9-mediated human plasmacytoid dendritic cell activation by interfering with IRF-7 and NF-κB signaling"

### Slide 1
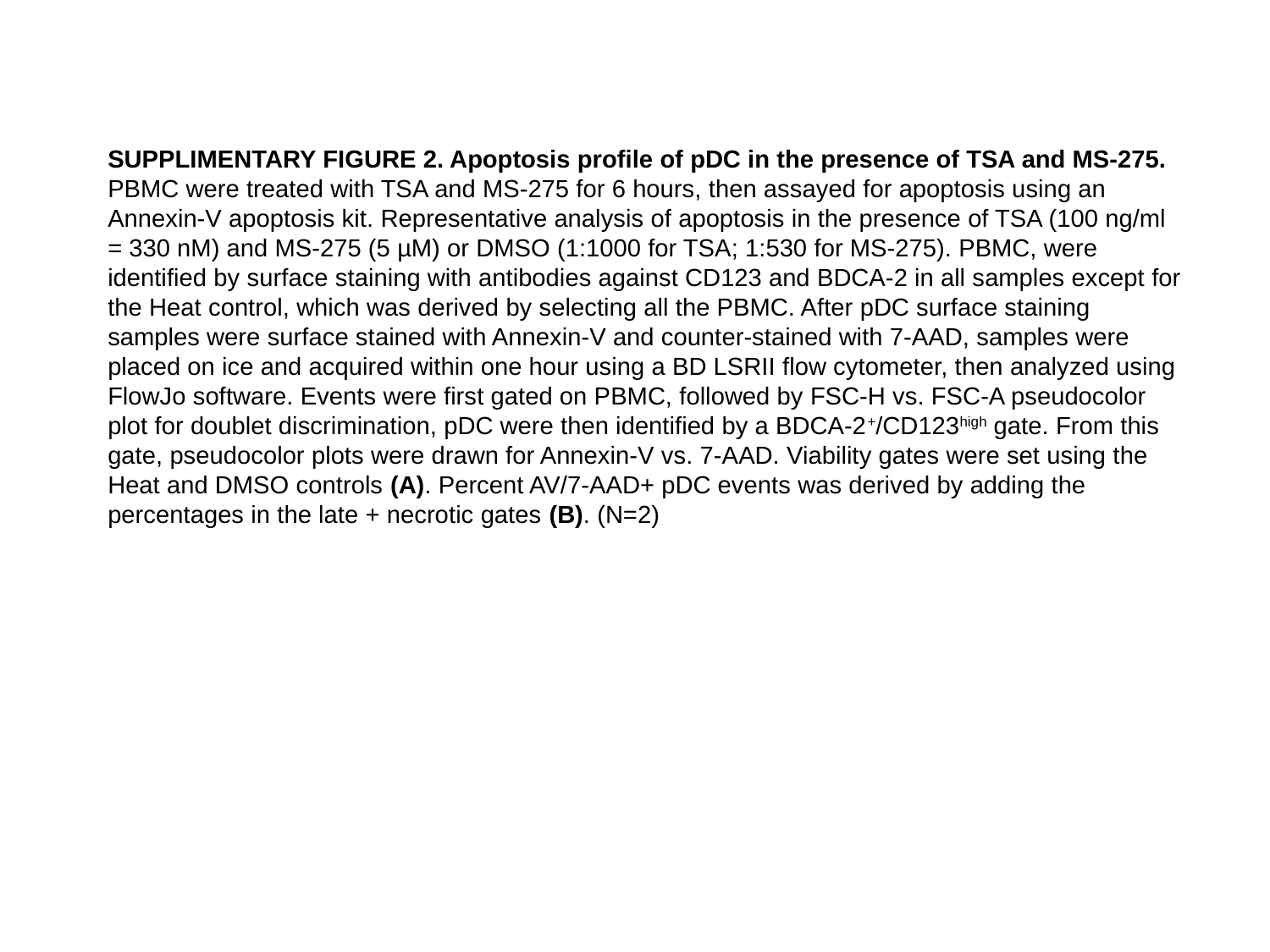

SUPPLIMENTARY FIGURE 2. Apoptosis profile of pDC in the presence of TSA and MS-275. PBMC were treated with TSA and MS-275 for 6 hours, then assayed for apoptosis using an Annexin-V apoptosis kit. Representative analysis of apoptosis in the presence of TSA (100 ng/ml = 330 nM) and MS-275 (5 µM) or DMSO (1:1000 for TSA; 1:530 for MS-275). PBMC, were identified by surface staining with antibodies against CD123 and BDCA-2 in all samples except for the Heat control, which was derived by selecting all the PBMC. After pDC surface staining samples were surface stained with Annexin-V and counter-stained with 7-AAD, samples were placed on ice and acquired within one hour using a BD LSRII flow cytometer, then analyzed using FlowJo software. Events were first gated on PBMC, followed by FSC-H vs. FSC-A pseudocolor plot for doublet discrimination, pDC were then identified by a BDCA-2^+^/CD123^high^ gate. From this gate, pseudocolor plots were drawn for Annexin-V vs. 7-AAD. Viability gates were set using the Heat and DMSO controls (A). Percent AV/7-AAD+ pDC events was derived by adding the percentages in the late + necrotic gates (B). (N=2)

### Slide 2
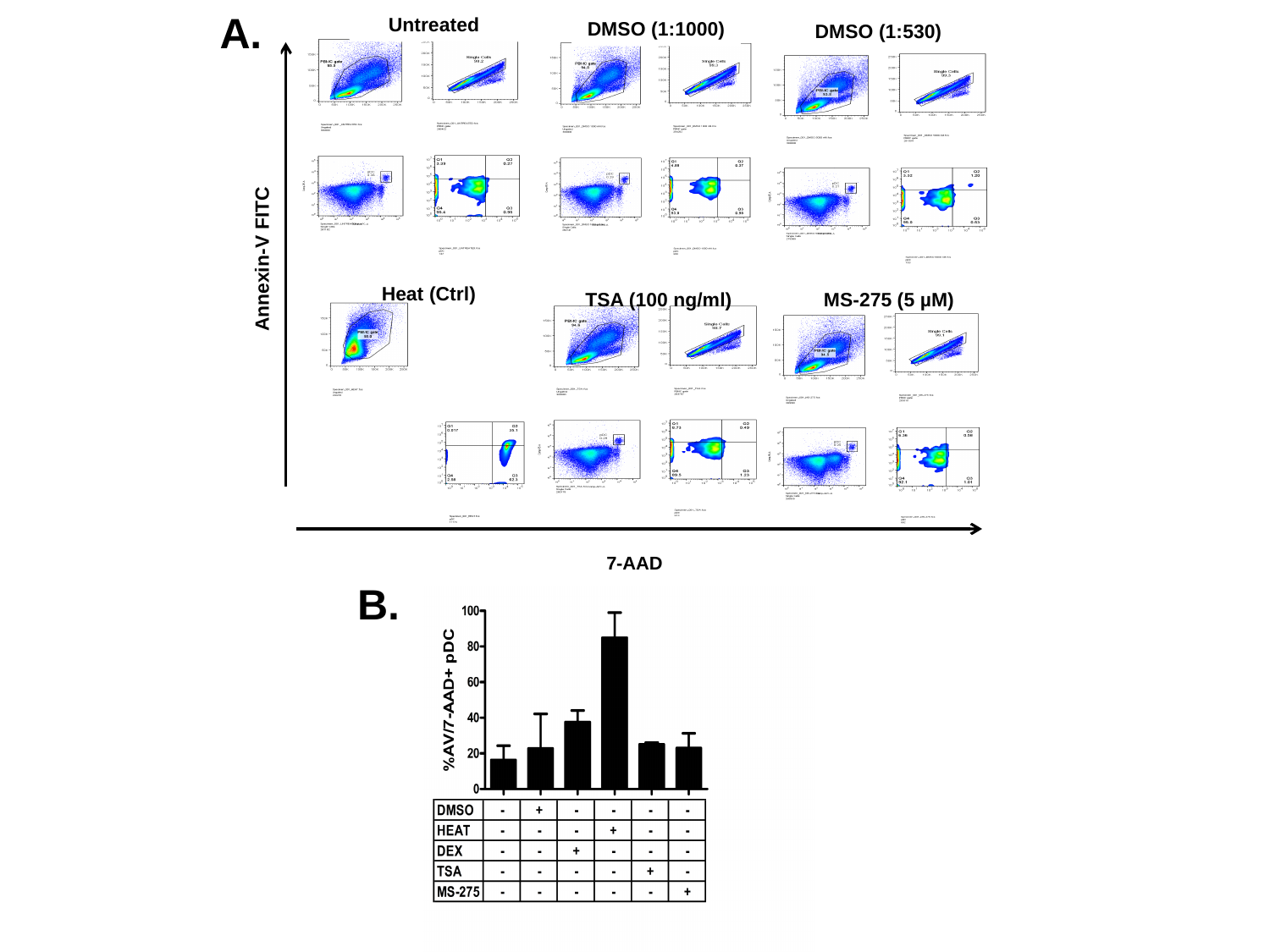

A.
Untreated
DMSO (1:1000)
DMSO (1:530)
Annexin-V FITC
Heat (Ctrl)
TSA (100 ng/ml)
MS-275 (5 µM)
7-AAD
B.
